# Divergent activation of the RXFP1 relaxin receptor by protein and small molecule agonists

**DOI:** 10.64898/2026.06.10.731386

**Authors:** James Osei-Owusu, Tristan Girbau, Xunan Wang, Jeffrey S. Smith, Pengxiang Shen, Sarah C. Erlandson, Amanda K. Williamson, Xiaojing Cong, Mark W. Grinstaff, Daniel E. Kahne, Cherine Bechara, Andrew C. Kruse

## Abstract

RXFP1 is an unusual G protein-coupled receptor (GPCR) that mediates physiological adaptations to pregnancy and is a therapeutic target due to its benefits in treatment of fibrosis and heart failure. Protein and small-molecule agonists are currently in mid-stage clinical trials. Here, we present three cryo-electron microscopy structures of RXFP1: ligand-free, bound to the native agonist relaxin-2, and bound to the small-molecule drug candidate AZD5462. These structures show that relaxin-2 engages the receptor’s ectodomain while AZD5462 binds within transmembrane domain. Together with hydrogen-deuterium exchange mass spectrometry, we show that relaxin-2 induces conformational reorganization of the linker domain into a helical secondary structure. In contrast, AZD5462 stabilizes a unique active conformation capable of inducing β-arrestin recruitment. Both mechanisms are distinctive from the “push-pull” mechanism of glycoprotein hormone receptors. Altogether, our findings define divergent activation mechanisms for protein and small-molecule agonists of RXFP1 and provide structural framework for next-generation drug discovery targeting relaxin receptors and their relatives.

## Introduction

The relaxin receptor RXFP1 is a member of the leucine-rich repeat-containing GPCR family (LGR), defined by a large N-terminal extracellular domain (ECD) made up of leucine-rich repeats (LRRs).^1^ RXFP1 and RXFP2 belong to a sub-family of the LGRs that also include a low-density lipoprotein receptor class A (LDLa) module and a linker domain.^1^ The LDLa module is required for receptor activation but is dispensable for ligand binding.^2,3^ RXFP1 and its cognate agonist, relaxin-2,^4^ serve as critical mediators of the myriad physiological changes that occur during pregnancy.^5^ Relaxin-2 is a small peptide hormone comprising two polypeptide subunits (A and B chains) connected by disulfide bonds. It belongs to the insulin-like superfamily of peptides.^6^ Although initially identified as a pregnancy hormone, relaxin-2 regulates RXFP1 signaling in renal and cardiovascular biology in both males and females.^4^ RXFP1 is an emerging therapeutic target for cardiovascular diseases, muscoskeletal fibrosis, kidney diseases, and cancers of reproductive tissues.^7–13^ Its activation by relaxin-2 induces several signaling pathways including cyclic adenosine monophosphate (cAMP) elevation that leads to increased cardiac output, increased renal blood flow, and modification of connective tissues.^14–16^ Notably, treatment with relaxin-2 specifically reverses cardiac fibrosis and arthrofibrosis in animal models, highlighting its antifibrotic potential.^13,17–19^

Given these and other data, substantial interest exists in RXFP1 as a therapeutic target. Recombinant relaxin-2 (Serelaxin) failed in a large multi-center Phase 3 clinical trial in patients with acute heart failure, a result attributed at least in part to the short circulating half-life of relaxin-2.^20–24^ At early treatment timepoints, significant benefits were seen in kidney function and mortality, prompting the development of next-generation long-acting relaxin receptor agonists.^20,23,24^ Several such molecules are now in phase 2 clinical trials, primarily for various forms of heart failure. For instance, AZD3427^25^ and TX45^26^ trials are currently on-going in stable patients with pulmonary hypertension in heart failure with preserved ejection fraction (PH-HFpEF), while volenrelaxin trial showed improved left atrial function at low dose but with worsened congestion in patients with worsening HFpEF.^27^ In addition to protein drug candidates, small molecules also offer a potential path to develop agonists targeting RXFP1 for chronic conditions.^28,29^ One of the most advanced is AZD5462, a recently developed RXFP1 small molecule agonist that is currently in Phase 2 clinical trial for chronic heart failure.^29,30^ Given the clear therapeutic potential and important native biology of RXFP1, understanding the details of this receptor’s activation and inhibition is of critical importance.

Previous studies using bioluminescence resonance energy transfer (BRET) and nuclear magnetic resonance (NMR) led to the hypothesis that relaxin-2 binds in the ECD, which causes conformational changes in the LDLa module and linker to trigger receptor activation, thus providing evidence for a multi-component molecular mechanism.^31,32^ We therefore set out to elucidate the basis of RXFP1 activation by providing atomic-level information through single particle cryo-electron microscopy (cryo-EM). We previously solved an active-state cryo-EM structure of the transmembrane domain (TMD) of RXFP1 and demonstrated that its extracellular loop 2 (ECL2) plays a key regulatory role in RXFP1 activation by occupying the orthosteric ligand-binding pocket in the active state, highlighting the critical role of the ECL2 in RXFP1 TMD activation.^33^ However, this progress is limited by a lack of structural data for the inactive state and an unclear understanding of how ligand binding induces receptor activation.

Here, we report the cryo-EM structures of the relaxin-2 receptor RXFP1 in its apo (ligand-free) and relaxin-2-bound states, as well as the cryo-EM structure of RXFP1-miniGs (minimal Gα protein) bound to the small-molecule drug candidate, AZD5462. While relaxin-2 binds to the ECD, AZD5462 binds to an unexpected site in the TMD on the lipid-facing side of transmembrane helix 7 (TM7), distinct from all other known GPCR ligand-binding sites.^34^ The identification of the AZD5462-binding pocket opens new avenues for designing small-drug molecules targeting RXFP1 and related receptors. Using a cAMP signaling assay, we also demonstrate that the TM7 and the hinge region are key regulators of RXFP1 activation and inhibition. Together, this study provides important structural and functional information for understanding the molecular mechanisms of RXFP1 activation by relaxin-2 and AZD5462, and for lead optimization and drug development.

## Results

### Structure determination of the ligand-free and relaxin-2-bound RXFP1

To elucidate the activation mechanism of RXFP1, we determined the cryo-EM structures of RXFP1 in its apo and relaxin-2-bound states (**Figure 1**). We had previously solved an active-state structure of the receptor bound to a relaxin-2 analog (SE001: single-chain relaxin-2) and proposed that RXFP1 activation occurs through a mechanism of autoinhibition. In this mechanism, the receptor’s extracellular loop 2 occupies the canonical GPCR orthosteric site in the active state but is inhibited by the ECD in the absence of relaxin-2.^33^ However, this previous work was limited by the structural uncoupling of the ECD from the TMD, resulting in poor resolution of the ECD region. Moreover, the lack of structural data for an inactive conformation of the receptor limited our ability to interpret structural details of receptor activation.

**Figure 1.**
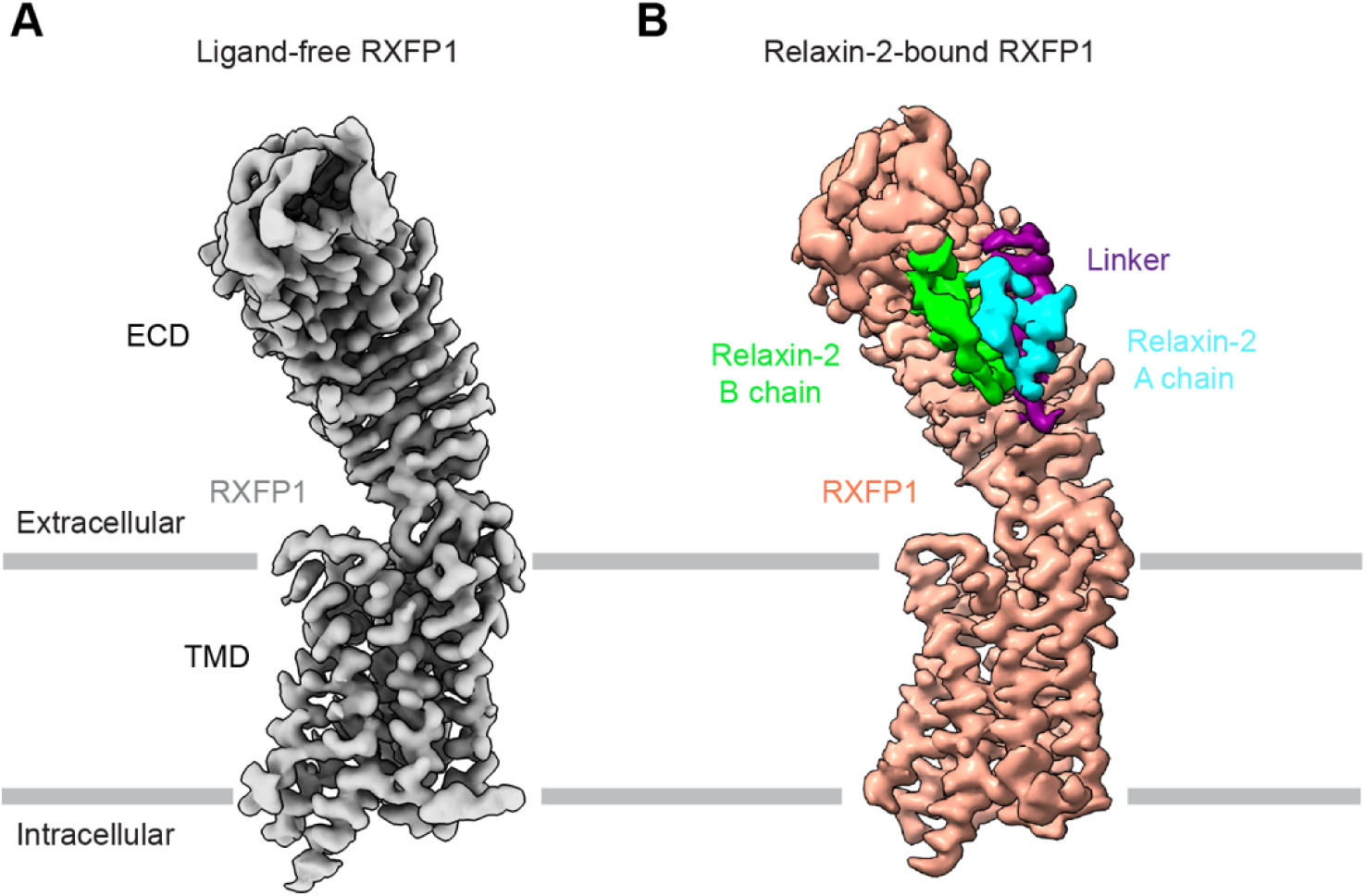
Cryo-EM maps of ligand-free and relaxin-2-bound RXFP1. (**A**) Cryo-EM map of ligand-free RXFP1 (EMD-75403), colored in gray. RXFP1 domains observed in the map are the transmembrane domain (TMD) and ectodomain (ECD – consisting of the leucine-rich repeats (LRR) and the hinge region). (**B**) Cryo-EM map of relaxin-2-bound RXFP1 (EMD-75473). RXFP1 TMD, hinge and LRR are colored orange, and RXFP1 linker domain is colored purple. Relaxin-2 A chain is colored cyan and relaxin-2 B chain is colored green.

To address these limitations, we cloned RXFP1 fused to the engineered Gα miniGs as we previously reported (**Figures S1A-S1C**).^33,35,36^ A 3C proteolysis site was inserted between RXFP1 and the miniGs to enable inactive-state structure determination (**Figures S1A-S1C**). This allowed miniGs cleavage and subsequent release of RXFP1 from the active state conformation (**Figures S1C and S1D**).^35,36^ The inactive-state structure of RXFP1 was determined by cryo-EM to a global resolution of 3.5 Å, with clear density for the TMD, hinge and LRR domains, allowing us to build an atomic model (**Figures 1A and S1H-S1M; Table S1**). In contrast, the densities for the linker and LDLa module were not observed, indicating flexibility of these regions.

To understand the recognition of relaxin-2 by RXFP1, we solved a relaxin-2-bound state structure of RXFP1 at a global resolution of 3.5 Å using the same method of proteolytic miniGs removal (**Figures 1B, S1F, S1G and S2A-S2E**). Unlike the ligand-free structure where the linker domain was not observed, the relaxin-2-bound state structure showed clear density for the linker domain, suggesting that the binding of relaxin-2 stabilized this region (**Figures 1, S2F and S2I**). The LDLa module, on the other hand, was not observed in this structure despite multiple efforts to obtain a map with density for the LDLa module. Both local refinements of our structures and attempts to stabilize RXFP1 using a G protein complex did not allow visualization of the LDLa module. Our final cryo-EM map was sufficient to position components of RXFP1 and the two subunits of the relaxin-2 hormone, the A chain and B chain (**Figures S2F-S2G; Table S1**). Structural analysis showed that the relaxin-2-bound structure is in an *inactive* conformation, characterized by an inward position of its transmembrane helix 6 (TM6) (**Figure S2H**). This result suggests that our relaxin-2-bound inactive-state structure may represent an intermediate step on the pathway of receptor activation, where hormone has bound but G protein is not yet engaged. This may also account for the lack of structural uncoupling between the ectodomain and transmembrane domain, which appears to be a feature unique to the fully activated state of the receptor.^33^ The incomplete activation of agonist-bound receptor is a common feature of agonist-bound but G protein-free GPCRs, since ligand binding alone is generally insufficient to fully activate the receptor.^37^

### Molecular Recognition of Relaxin-2 by RXFP1

Our structures show that the ECD of RXFP1 contains 14 LRRs and forms an arc-shaped tube with parallel beta-sheets forming the concave side, similar to glycoprotein hormone receptors (GPHRs) and toll-like receptors (TLRs) (**Figures 1 and 2A**).^38–40^ Relaxin-2 binds to the concave side, spatially separated from the TMD and not making direct contact, contrary to previous proposals of a secondary binding site for relaxin-2 on the ECLs.^41^ The interactions of relaxin-2 and RXFP1 are mediated by both A and B chain subunits of relaxin-2, with the B chain directly interacting with the LRR and the A chain interacting with the linker (**Figure 2**). This is consistent with the current model of RXFP1 activation that proposes that both A and B chains are involved in binding to the ECD, and that the B chain is the primary site for interaction.^31,32,42,43^ The two chains of relaxin-2 are connected by two disulfide bonds at C11^B^-C11^A^ and C23^B^-C24^A^ (**Figures 2A and S2G**). In addition, the A and B chains are connected by hydrogen bonds including K17^A^-E6^B^ and Y3^A^-Q19^B^-L20^A^ (**Figure S2G**). The C-terminal half of the B chain (R13^B^-T27^B^) forms a helix that packs against the LRR region whereas the N-terminal half (S2^B^-G12^B^) makes direct contact with the A chain (**Figures 2A and S2G**). The surface of B chain relaxin-2 contains positively charged residues, which form complementary electrostatic interactions with negatively charged residues on the surface of RXFP1 ECD (**Figures 2C and 2D**). Specifically, R17^B^ and R13^B^ protrude from the B chain helix, each directly interacting with a pair of acidic residues on the RXFP1 LRR, D253/E255 and E299/D301 respectively (**Figure 2C**). Mutating these residues to alanine destabilizes the interactions and hence significantly reduced relaxin-2 binding and consequently RXFP1 activation, suggesting that these residues are key for relaxin-2 recognition of RXFP1 (**Figure 2F; Table S2**). These results are concordant with the previously reported loss of activity in mutations of R17^B^ and R13^B^.^44–46^ Accordingly, our results confirm prior studies that identified D253/E255 and E299/D301 as critical binding residues in RXFP1,^42^ but provide key structural context to their role in RXFP1 activation.

**Figure 2.**
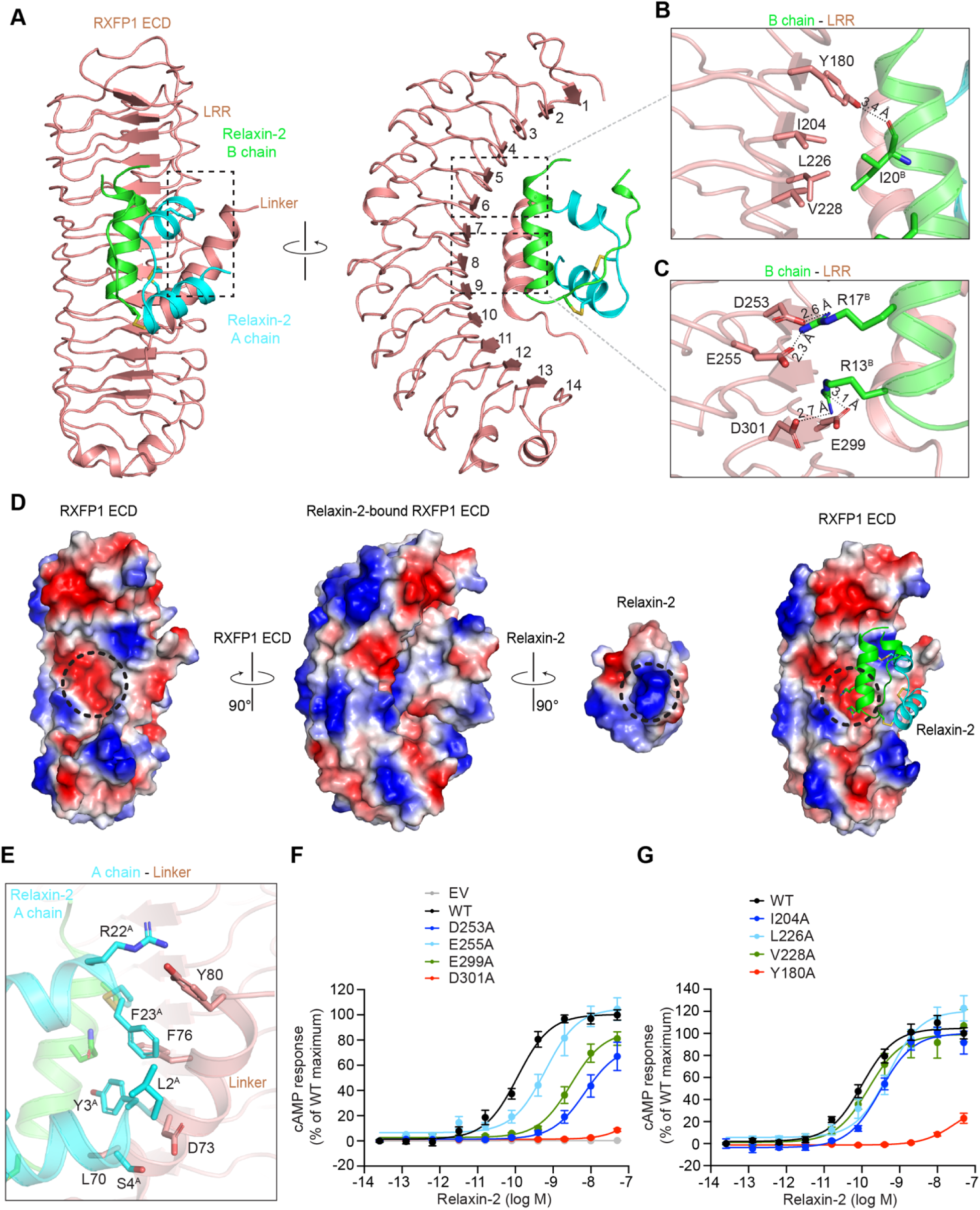
Interactions between Relaxin-2 and RXFP1. (**A**) The interface of RXFP1 ECD (brown) and relaxin-2 hormone, A chain (cyan) and B chain (green). (**B-C**) The interface of RXFP1 LRR and the B chain of relaxin-2. (**D**) Surface charge distribution of relaxin-2 and RXFP1 ECD only, relaxin-2-bound RXFP1 ECD, relaxin-2 only and relaxin-2 (model)-bound RXFP1 ECD. Dashed circle represents region of charged residues interaction between the ECD and relaxin-2. (**E**) The interface of RXFP1 linker and A chain of relaxin-2. (**F-G**) cAMP concentration-response curves for point mutations of residues in RXFP1 LRR interacting with relaxin-2 B chain subunit. For cAMP response, 100% is the maximum activity of the WT receptor. The pEC_50_, half-maximal effect, and maximum response are outlined in Table S2. Data are mean ± s.e.m. from at least three independent experiments.

A previously predicted hydrophobic interaction between I20^B^ and a cluster of non-polar residues, I204, L226 and V228 was also observed (**Figure 2B**).^42^ Surprisingly, mutations of these RXFP1 residues to alanine did not significantly affect receptor signaling (**Figure 2F**). Instead, the carbonyl group of I20^B^ forms a hydrogen bond with Y180 and loss of function mutation of Y180 resulted in reduced receptor activity (**Figure 2G; Table S3**). In contrast to the B chain, the A chain does not directly interact with the LRR domain. Rather, it makes close contact with the linker domain through hydrophobic interactions between its N- and C-termini and the linker (**Figures 2A and 2E**). This is consistent with prior data that showed that the linker contained a site for relaxin-2 binding and proposed that a combination of this site and the LRR binding site are both required for the sub nanomolar affinity of relaxin-2 for RXFP1.^32^ Previous data has also shown that the linker domain is important for relaxin-2 binding and RXFP1 activation,^31–33,47,48^ however, mutating individual residues involved in the hydrophobic interactions did not significantly affect receptor signaling (**Table S3**), suggesting that residues in this region work collectively to promote receptor activation.^49^ This finding, in addition to the complete loss of relaxin-2 signaling associated with the B chain-LRR mutations provide further support to previous studies that the B chain is the primary site of relaxin-2-RXFP1 interaction.

Binding of relaxin-2 to the ECD stabilized a portion of the linker domain (N66-Y80) into a helical structure that packs against the concave side of the LRR (**Figures 2A and S2K**).^32^ Interestingly, residues D73-Y80 of the linker helix are absent in RXFP2. This might explain why relaxin-2 has a much lower affinity for RXFP2.^47,48^ Residues of the linker connecting the helix to the LRR were not observed in our structure. Similarly, the LDLa domain was not observed, suggesting flexibility in these regions. In order to measure relaxin-2-induced structural changes in RXFP1 and gain more insight into the overall structural arrangement of RXFP1, we employed hydrogen-deuterium exchange mass spectrometry (HDX-MS) and compared the deuterium exchange for ligand-free and relaxin-2-bound RXFP1. Although we observed limited coverage of the receptor transmembrane region, as is typical with membrane proteins, a significant decrease in deuterium uptake in regions of the LDLa module, D58-C62, and the linker domain, G63-F72, occurs in the relaxin-2 bound state, indicating that conformational changes shield the backbone amides and hence occlude access to deuterium (**Figures S2I, S2J and S2L**). This decreased uptake in the presence of relaxin-2 was most pronounced at early deuteration time points and gradually diminished with longer labeling times, suggesting that relaxin-2 induces local stabilization and dynamic structuring of this segment, which appears to remain largely unstructured in the ligand-free RXFP1. These data are consistent with our relaxin-2-bound cryo-EM structure that reveals density for the linker domain and shows that the linker adopts a defined secondary structure and directly interacts with the LRR (**Figures S2K and S2M**). This interaction is through a network of residues, including C62, D64, N66, W68, Q71, F72 and Y75 of the linker domain (**Figure S2M; Table S4**). Mutagenesis of these residues to alanine revealed D64 and W68 as important players in RXFP1 activation (**Figure S2N**). The LDLa module plays an important role in signal transduction during receptor activation even though our cryo-EM structures and HDX studies revealed little to no structural information. We propose that the reorganization of the linker domain into a helix on the LRR and occlusion of residues in the LDLa module upon relaxin-2 binding indicate that the LDLa module makes contact with the ECD and possibly with ECL1 and the TMD. These conformational changes associated with relaxin-2 binding may rearrange and stabilize the LDLa module, thereby increasing the local concentration of the LDLa module to mediate receptor activation. Altogether, our data shows that relaxin-2 binds to the ECD of the receptor including distinct interactions with both the LRR and the linker domains. Binding of relaxin-2 causes conformational changes in the linker and consequently the LDLa module to transduce activating signal from the ECD to the TMD.

### Autoinhibitory role of the hinge region in RXFP1 activation

Structural comparison of the inactive and the active states of GPHRs (belong to the same LGRs family as the relaxin receptors) such as luteinizing hormone/choriogonadotropin receptor (LHCGR), thyrotropin receptor (TSHR) and follicle-stimulating hormone receptor (FSHR), has revealed that the activation mechanism of the GPHRs involves major conformational movement of the entire LRR domain relative to the membrane layer upon ligand binding, generally referred to as the “push and pull” mechanism.^38–40,50^ Thus, we determined whether RXFP1 undergoes a similar conformational change upon relaxin-2 binding. The density for the ECD of the previously solved active-state RXFP1^33^ was poorly defined and did not allow for an atomic model building. However, we could still place the receptor LRR to compare with its position in the inactive-state structure (**Figures 3A and S3A**). To our surprise, we observed only 3.7° rotational shift of the LRR upon ligand binding (**Figure 3A**), in contrast to the drastic rotational shifts of 45° and 55° for LHCGR and TSHR respectively.^38,50^ This result is consistent with the conformational changes observed in the relaxin-2-bound inactive-state RXFP1, where we observed stabilization of the linker domain, but no major overall rotational shift in the LRR domain relative to the membrane layer (**Figures S3B and S3C**), further attesting to the importance of the conformational change associated with the linker domain in RXFP1 activation mechanism. A possible explanation for this difference is the size of the respective ligands. Relaxin-2 is substantially smaller in size at 6 kDa, compared to the glycoprotein hormones ranging from ∼28 to 46 kDa,^38,51–53^ and hence binding of relaxin-2 is unable to drastically push the ECD relative to the membrane layer as observed with the GPHRs. The stability of the RXFP1 ECD orientation on the TMD is further supported by a disulfide bond between C380 on the hinge helix and C389 on the LRR C-terminus (**Figures S3C and S3D**).

**Figure 3.**
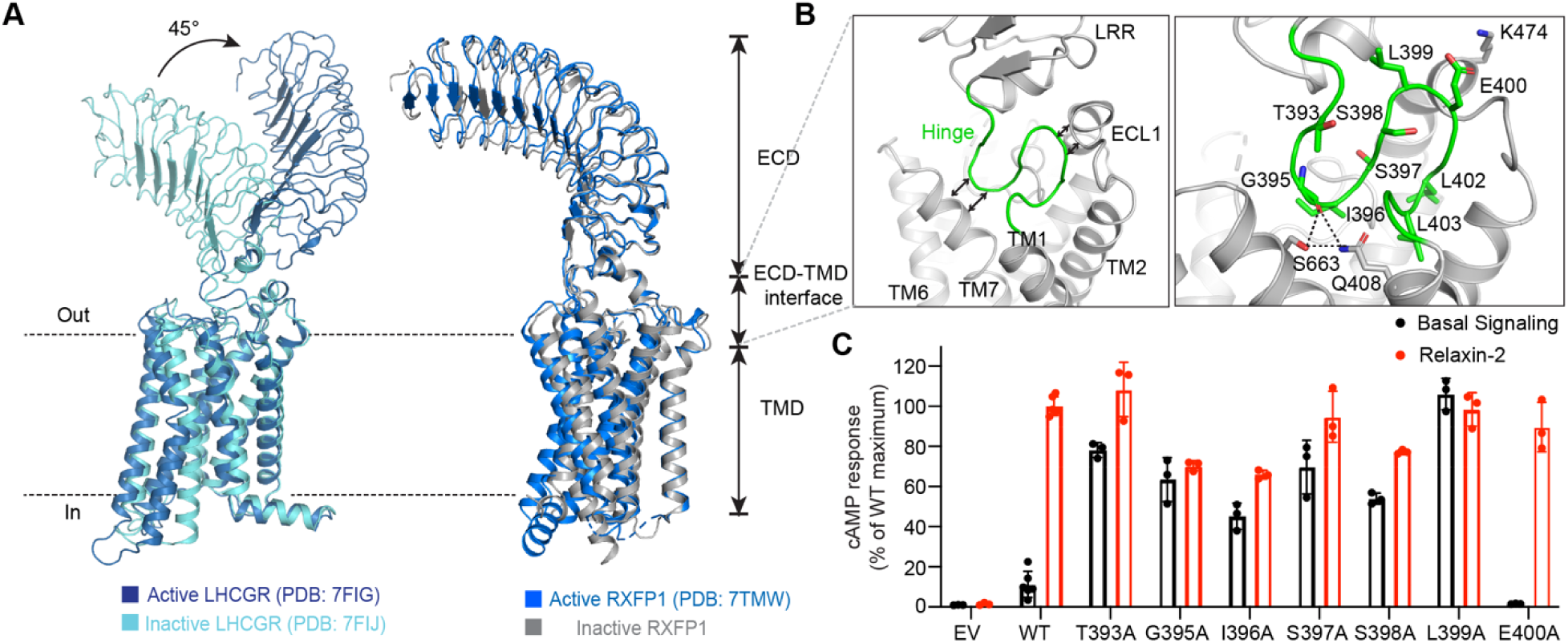
The hinge forms the ECD-TMD interface and stabilizes the inactive conformation. (**A**) Structural comparison of active and inactive LHCGR and RXFP1, with the TMD aligned. Active LHCGR (PDB: 7FIG, blue), Inactive LHCGR (PDB: 7FIJ, cyan), Active RXFP1 (PDB: 7TMW, blue) and Inactive RXFP1. (**B)** The hinge forms a loop (green) that is in close proximity to TM7 and ECL1. Details of ECD-TMD interface and interactions between the hinge region and TM7 residues. (**C**) Basal signaling and signaling in response to 10 nM relaxin-2 for hinge region mutants. Data are mean ± s.e.m (n = 3 biological replicates).

Next, we investigated how the binding of relaxin-2 to the distal ECD leads to activation in the TMD. The ECD connects to the TMD by a hinge region, forming an ECD-TMD interface, just like the hinge in the GPHRs (**Figures 3A and 3B**). The role of the hinge has been explored in GPHRs as a critical regulator of receptor signaling and has been shown to adopt different conformations in the active and inactive states of these receptors.^38–40,50^ Our structures revealed that the hinge region is 15 amino acids long (P391-S405) and forms a loop with two turns, one end of the turn extending and packing closely towards TM7 and the other turn towards ECL1 to stabilize the ECD (**Figure 3B**). Residues in the hinge region also make close contacts with ECL2. Due to these domain contacts and the importance of the hinge in GPHRs signaling, we hypothesized that the hinge loop may play a key regulatory role in structural communication from the ECD to the TMD in relaxin receptors. To investigate this, we performed an alanine mutagenesis scan of residues in the hinge domain of RXFP1. Interestingly, alanine mutants of residues making up the half of the hinge loop that extends towards TM7 resulted in constitutive activity of the receptor (**Figure 3C; Table S5**), implying an autoinhibitory role of the hinge in receptor activation. This unexpected role of the hinge region in RXFP1 activation implies that the hinge prevents the continuous activation of the receptor in the absence of relaxin-2 but is disrupted upon relaxin-2 binding, releasing the autoinhibition for receptor activation. Just like the GPHRs, the functional role of RXFP1 hinge region might be associated with conformational changes. The TM7 is in close proximity to the hinge in the inactive state and undergoes an inward movement in the active state, likely clashing with the hinge domain (**Figure 4A**). Also, structural comparison of the ligand-free and relaxin-2-bound inactive-state RXFP1 shows conformational changes in the hinge region, highlighting a possible conformational change of the hinge region in the active state (**Figures S3C and S3E**). Residues involved in this autoinhibition are conserved in RXFP2, suggesting a conserved autoinhibitory role of the hinge in relaxin receptors.

**Figure 4.**
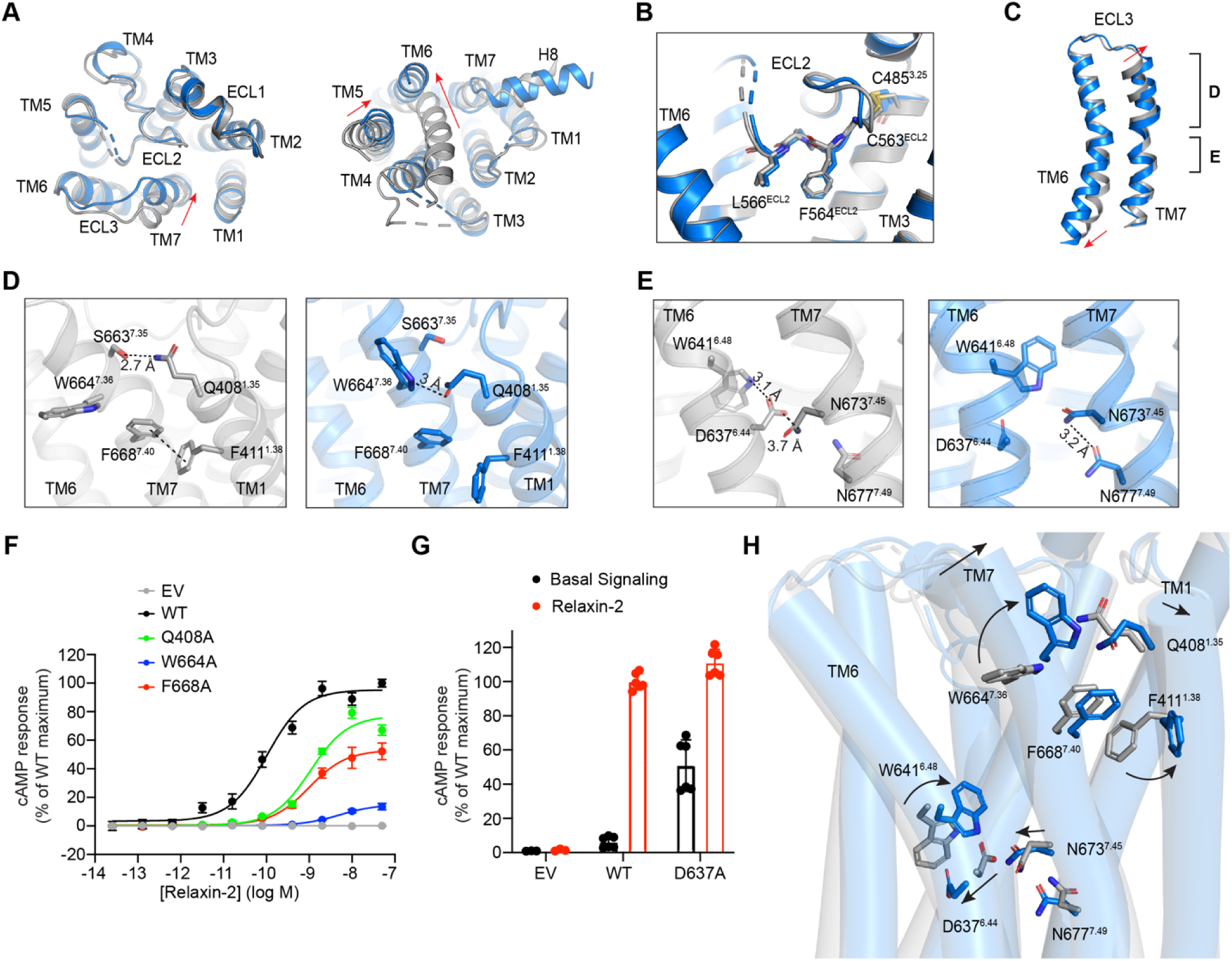
Activation mechanism of RXFP1 by relaxin-2. (**A**) Structural comparison of the active (blue) and inactive (grey) RXFP1, extracellular and intracellular views. (**B**) The conformation of ECL2 in active- and inactive-state RXFP1. Details of ECL2 in the canonical GPCR orthosteric ligand-binding pocket. (**C**) Structural comparison of TM6 and TM7 of active and inactive RXFP1. (**D**) Detailed interactions occurring on the upper region of TM7 and TM1, active and inactive RXFP1. W664 forms a hydrogen bond with Q408 in the active state. (**E**) Detailed interactions occurring on the mid-region of TM7 and TM6. The hydrogen bond between W641 and D637 is broken in the active state. (**F**) cAMP concentration-response curves for point mutations of residues in RXFP1 TM7 and TM1 in response to relaxin-2. Data are mean ± s.e.m. from at least two independent experiments. (**G**) Basal signaling and signaling in response to 10 nM relaxin-2 for D637A mutant. Data are mean ± s.e.m. from two independent experiments. (**H**) Overlay of active (blue) and inactive (grey) RXFP1 models, illustrating a stepwise mechanism of RXFP1 activation in the TMD. Sticks and arrows are shown for residues involved in conformational change upon activation by relaxin-2.

### Relaxin-2-mediated RXFP1-TMD activation

The TMD of our ligand-free inactive-state structure displays hallmarks of inactive-state GPCRs (**Figures 4A and S4A**). Most notably, TM6 is in the inward conformation, precluding G protein coupling. Additionally, we observed an outward shift of TM5 in the intracellular side and an outward movement of TM7 in the extracellular side, resulting in a conformational shift in ICL3 and ECL3 respectively (**Figure 4A**). A key unusual feature observed in the active-state structure was ECL2 occupying the canonical GPCR orthosteric ligand-binding pocket (**Figure 4B**). Residues F564^ECL2^ and L566^ECL2^ in ECL2 fit deep into the pocket and were shown to be essential for receptor activation.^33^ In comparing the conformations of ECL2 in the active and inactive-state receptor structures, we see minimal conformational difference for F564^ECL2^ and L566^ECL2^ (**Figure 4B**). However, nearby residues are rearranged as a result of the inward movement of TM5 and TM7, increasing hydrophobic contacts surrounding the orthosteric pocket in the active state compared to that of the inactive-state structure (**Figure S4B**). In light of our recent findings, we conclude that ECL2 acts a key signal transducer to mediate relaxin-2-dependent RXFP1 activation. This regulatory role of ECL2 in relaxin-2-mediated activation of RXFP1 is further highlighted by the rescue of activity of F564A^ECL2^ and L566A^ECL2^ mutants when treated with the small molecule agonist, AZD5462, implying that these residues are strictly required for relaxin-2 signaling but dispensable for AZD5462 signaling (**Figure S4C; Table S4**). The ECL2 of TSHR and LHCGR also occupies the canonical GPCR orthosteric ligand-binding pocket, however,^38,39^ mutagenesis of their equivalent residues did not affect receptor activation, further indicating the critical role of ECL2 in RXFP1 activation (**Figure S4D; Table S6**). It is important to note that the regions of ECL2 that face the extracellular domain, S568^ECL2^-S573^ECL2^, did not have clear density in either the active or inactive structures, suggesting intrinsic flexibility of this region in both states.

The most significant conformational change observed in the TMD of the active-state structure was the inward movement of the extracellular side of TM7 (**Figure 4A and 4C**). This conformational change plays a key role in receptor activation by triggering a network of residues along the TM6 and TM7 helices, notable among them are D637^6.44^, W641^6.48^, W664^7.36^ and F668^7.40^ (superscript indicates the Ballesteros-Weinstein numbering system). In the active state, W664^7.36^ is flipped upwards to form a hydrogen bond with Q408^1.35^ (**Figures 4D and 4H**). This stabilizes the inward position of the TM7 which breaks a π-π interaction between F411^1.38^ and F668^7.40^ (**Figure 4D and 4H**). Mutating Q408^1.35^, W664^7.36^ or F668^7.40^ to alanine significantly reduced or abrogated receptor activation (**Figure 4F; Table S3**), suggesting that these residues are important for RXFP1 activation. This inward movement of TM7 also consequently breaks a key hydrogen bond interaction between D637^6.44^ and W641^7.36^, that stabilizes the inactive state (**Figure 4E and 4H**). The D637^6.44^ and W641^7.36^ interaction is also stabilized by N673^7.45^, but this interaction is broken by the inward movement of TM7. Consequently, the sidechain of N673^7.45^ clashes with D637^6.44^, thereby disrupting the inactive conformation and causing the hallmark outward movement of TM6, a characteristic feature of GPCR activation (**Figure 4E**). This result is confirmed by the constitutive activity of the D637A mutant, highlighting its importance in stabilizing the inactive-state conformation of the receptor (**Figure 4G; Table S5**).

Our study show that both the hinge region and TM7 are key regulators of receptor activation, with the hinge region stabilizing the inactive-state conformation and the TM7 stabilizing the active-state conformation. Our results therefore provides structural and functional data that confirm that RXFP1 activation is indeed a multi-component process^31,32^ and further define a general model for relaxin-2-mediated RXFP1 activation; 1) the binding of relaxin-2 to the LRR and linker domains of RXFP1, with extensive polar interactions between the B chain of relaxin-2 and the LRR, and low affinity binding between the A chain and the linker, 2) rearrangement of the linker into a helical secondary structure to facilitate close contacts between the linker and the LRR, and consequently influence the conformation and interactions of the LDLa module, 3) stabilization of the hinge region into an active conformation, 4) conformational changes in the TMD that increase the hydrophobic contacts around ECL2, and, 5) conformational changes in TM7 and consequently TM6 to drive receptor activation (**Figure 7**).

### The small molecule AZD5462 uniquely activates RXFP1 and recruits β-arrestin

Discovering small molecules that activate RXFP1 has proven to be difficult despite considerable effort.^28^ Until recently, only one published series of small molecule agonists existed, with a potency of around 100 nM for the best molecule, ML290,^28,54,55^ making it insufficient for clinical use. Optimization of ML290 led to the identification of a clinical candidate, AZD5462 (**Figure 5A**).^29^ AZD5462 activates RXFP1 in the absence of relaxin-2 (**Figure 5B**) and is the only RXFP1-targeting small-molecule candidate that has reached mid-stage clinical trials, highlighting its relevance in drug discovery for RXFP1. We therefore sought to investigate how this highly divergent agonist activates RXFP1 in comparison to relaxin-2.

**Figure 5.**
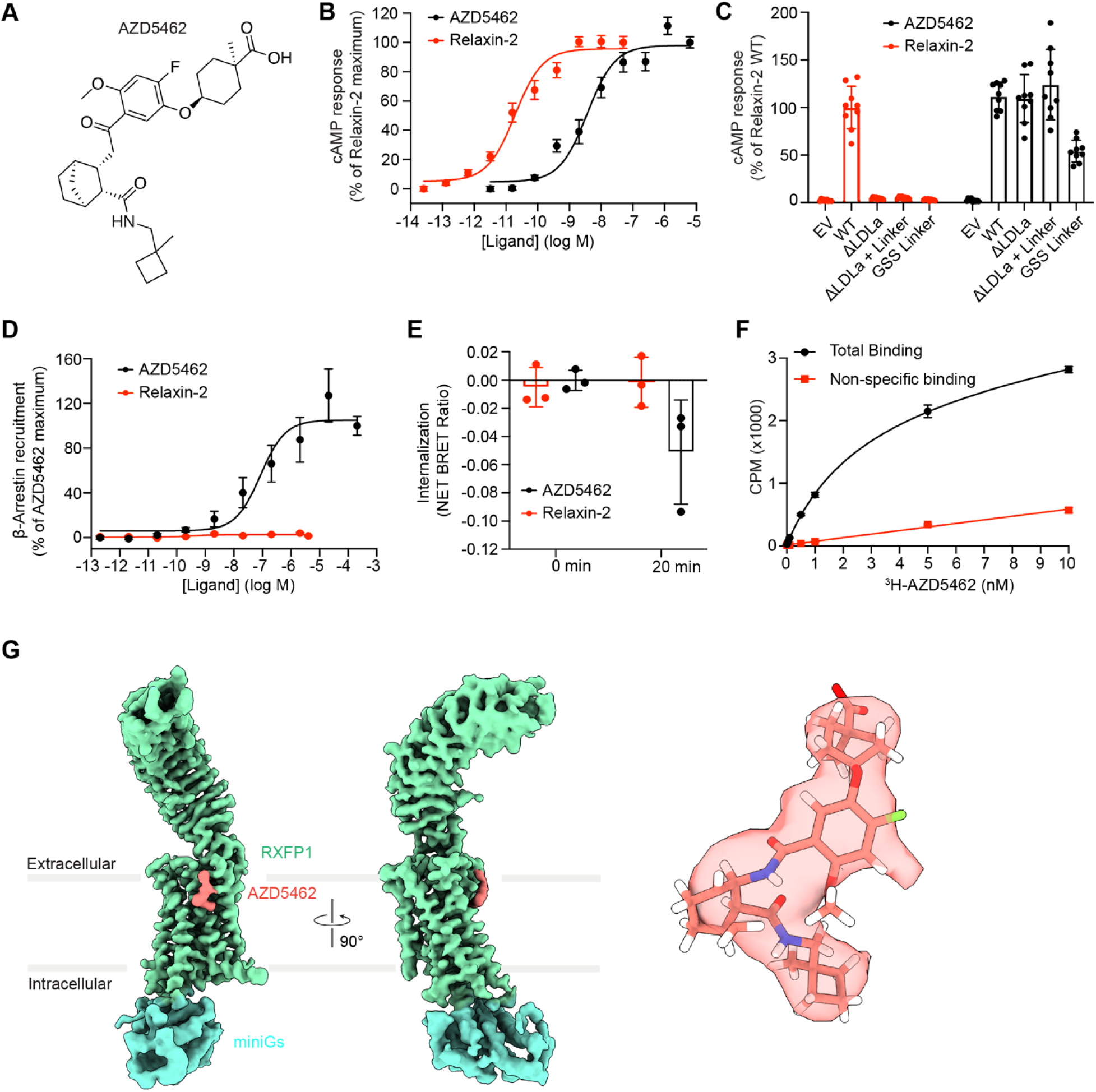
Functional and structural overview of AZD5462 activation of RXFP1. (**A**) Chemical structure of the RXFP1 agonist AZD5462. (**B**) cAMP concentration-response curves for WT RXFP1 in response to relaxin-2 or AZD5462. Data are mean ± s.e.m. from at least three independent experiments. (**C**) Signaling in response to 10 nM relaxin-2 or 1.25 μM AZD5462 for RXFP1 ECD mutants. Data are mean ± s.e.m. from three independent experiments. (**D**) β-arrestin concentration-response curves for WT RXFP1 in response to relaxin-2 or AZD5462. Data are mean ± s.e.m. from at least three independent experiments. (**E**) AZD5462 (10 μM) or relaxin-2 (0.1 μM)-mediated RXFP1 internalization for 0- or 20-min. Pre-read BRET ratios were subtracted from post-read BRET ratios, and these values were normalized to vehicle treated wells to obtain the NET BRET ratio. Data are represented as the means ± s.e.m. of three independent experiments. (**F**) Total and non-specific binding of radiolabeled AZD5462 (^3^H-AZD5462) to cell membranes containing RXFP1. Data are mean ± s.e.m. from three independent measurements. (**G**) Cryo-EM map of AZD5462-bound RXFP1-miniGs complex (EMD-75439), RXFP1 (green), miniGs (cyan) and AZD5462 (red). The structure and cryo-EM density of AZD5462 in the bound structure.

The LDLa module and the linker domain have long been known to be required for RXFP1 activation by relaxin-2. To test whether these features are also necessary for AZD5462-mediated activation, we cloned constructs of RXFP1 with deletion of the LDLa module, deletion of the LDLa module and linker domain, or substitution of the linker residues with GlyGlySer linker, and in each case, we tested RXFP1 activation using either relaxin-2 or AZD5462. Indeed, unlike relaxin-2-induced activation, AZD5462-mediated activation of RXFP1 does not depend on the LDLa module or the linker domain (**Figure 5C**), suggesting that AZD5462 and relaxin-2 might activate RXFP1 using different mechanisms. This result is consistent with our HDX experiments, where we compared the deuterium exchange for ligand-free and AZD5462-bound RXFP1 and observed no significant decrease in deuterium uptake in the linker domain and LDLa module (**Figure S2I and S2L**). Similarly, LRR residues, particularly Y180, D253 and D301, previously identified to be critical for relaxin-2-mediated activation of RXFP1 were dispensable for AZD5462-mediated activation, further highlighting a different mechanism of RXFP1 activation (**Figure S5A**).

Next, we tested the ability of AZD5462 to recruit β-arrestin using the PRESTO-TANGO system in HTLA cells overexpressing RXFP1. Previous studies have shown that relaxin-2 activation of RXFP1 does not recruit β-arrestin and exhibits poor internalization.^56^ Consistently, the PRESTO-TANGO assay revealed that relaxin-2 is unable to recruit β-arrestin, however, to our surprise, AZD5462 recruited β-arrestin to RXFP1 (**Figure 5D**). β-arrestins regulate cellular signaling by desensitization and subsequent internalization of receptors. We therefore tested RXFP1 internalization using a BRET internalization assay. AZD5462 induced internalization of RXFP1 whereas no significant internalization was observed for relaxin-2-mediated activation of RXFP1 (**Figures 5E and S5B**). These results indicate divergent signaling profiles of relaxin-2 and AZD5462, with only the latter inducing arrestin recruitment.

### Molecular recognition and activation of RXFP1 by AZD5462

Next, we determined the molecular basis by which AZD5462 activates RXFP1. Using ^3^H-labeled version of AZD5462, we measured the affinity of AZD5462 to RXFP1. AZD5462 binds to RXFP1-miniGs with a K_D_ value of ∼3 nM (**Figure 5F**). We then determined the cryo-EM structure of RXFP1-miniGs bound to AZD5462 (**Figures 5G and S5C-S5H**). The cryo-EM map was sufficiently clear to dock the AZD5462 molecule and build an atomic model for the AZD5462-RXFP1 complex (**Figures 5G and S5I**). Like the ligand-free inactive-state structure, densities for the LDLa module and linker were not observed. The AZD5462-bound RXFP1 structure represents an active-state structure, characterized by an outward position of its TM6 (**Figure S5K**) and G protein αs subunit bound to the receptor’s intracellular face (**Figure S5I)**. No major conformational differences in the position of LRR domain relative to our other active and inactive structures was observed (**Figure S5J**). AZD5462 binds to an unusual site of the receptor, far from the relaxin-2 binding site at the top half of TM7 and TM1, on the lipid-facing side of the helices (**Figures 5G and S5I**). This explains the non-essentiality of the LDLa module and linker for RXFP1 activation by AZD5462. To the best of our knowledge, this is the first observation of a small molecule binding to this location of a class A GPCR.^34^

Given that AZD5462 and ML290 share similar molecular architectures,^29^ we hypothesized that ML290 binds to the same pocket to activate RXFP1. To test this, we performed a radioligand competition assay using radiolabeled AZD5462 and cold ML290. Indeed, our assay showed that ML290 competes with AZD5462 for the same ligand-binding site (**Figure S5L)**, implying that both ML290 and AZD5462 bind to the same membrane-facing pocket to activate RXFP1.

The region of TM7 that serves as a binding pocket for AZD5462 is the same site that undergoes an inward movement in the single-chain relaxin-2-bound active-state structure, consistent with the role of TM7 in RXFP1 activation (**Figures 4 and 5G**). However, unlike the single-chain relaxin-2 active state, TM7 in the AZD5462-bound structure does not fully undergo this inward movement, suggesting that the inward collapse of TM7 is not required for AZD5462-mediated activation of the receptor (**Figure 6H**). AZD5462 binding to RXFP1 involves several residues, some of which were identified as key residues in the relaxin-2-mediated activation of the receptor. The interactions include hydrogen bonds between the head carboxylic group of AZD5462 and S405^1.32^ and Q408^1.35^, the middle carbamoyl group and W664^7.36^ and a π-π bond between the middle phenoxy ring and the aromatic ring of F668^7.40^ (**Figures 6A and 6B**). In addition, AZD5462 forms extensive hydrophobic interactions with the residues along the TM7 and TM1 helices, specifically between the middle 4-methoxy-phenyl group and V415^1.42^, the bridged cyclohexane group and I672^7.44^ and the tail 1-methylcyclobutyl group and P671^7.43^ (**Figures 6A and 6B**). We assessed AZD5462-mediated activation of WT and mutant RXFP1 using cAMP accumulation and β-arrestin recruitment assays. Alanine mutations of W664^7.36^ and F668^7.40^ completely ablated receptor cAMP activity and β-arrestin recruitment even though receptor expression was unaffected, indicating that these residues play important roles in transmitting AZD5462-mediated RXFP1 activation signal (**Figures 6C and 6D; Table S7**). Both W664^7.36^ and F668^7.40^ were identified as key players in relaxin-2-mediated activation of RXFP1. Relaxin-2 and AZD5462-mediated activity were completely ablated in W664A whereas only AZD5462-mediated activity was completely abolished in F668A, showing that F668^7.40^ is strictly required for AZD5462-mediated activity but not for relaxin-2-mediated activity (**Figure 6G; Table S4**).

**Figure 6.**
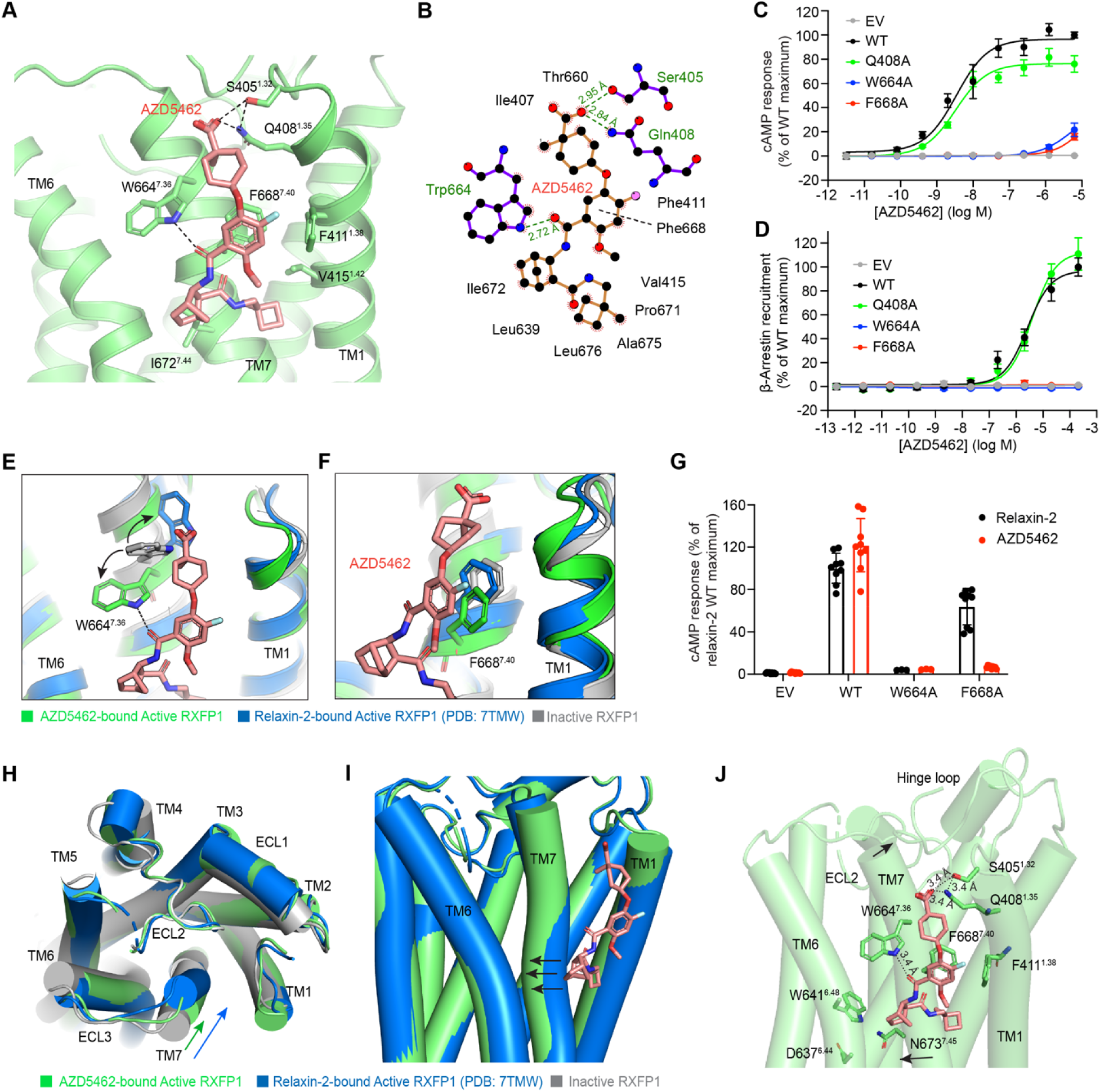
Activation mechanism of RXFP1 by AZD5462. (**A**) Detailed interactions between AZD5462 and RXFP1. (**B**) Schematic representation of AZD5462–RXFP1 interactions generated using ligPlus program.^58^ (**C**) cAMP concentration-response curves for point mutations in TM7 and TM1 in response AZD5462. Data are mean ± s.e.m. from at least three independent experiments. (**D**) β-arrestin concentration-response curves for point mutations in TM7 and TM1 in response AZD5462. Data are mean ± s.e.m. from at least three independent experiments. (**E-F**) Conformational changes of **E**) W664 and **F**) F668 in the ligand-free inactive (grey), relaxin-2 active (blue) and AZD5462 (green) active RXFP1. (**G**) Signaling in response to 10 nM relaxin-2 (black) or 1.25 μM AZD5462 (red) for RXFP1 TM7 mutants, W664A and F668A. Data are mean ± s.e.m. from 3 independent experiments. (**H**) Conformational comparison between ligand-free inactive (grey), relaxin-2 (blue) active and AZD5462 (green) active states. (**I**) Demonstrating the mid-region of TM7 inward push of the bulky leg of AZD5462. (**J**) Model of RXFP1 activation by AZD5462.

We next performed a structural comparison between the AZD5462-bound active-state, single-chain relaxin-2-bound active-state, and ligand-free inactive-state structures of RXFP1 and found conformational changes in these residues that contribute to their role in receptor activation. W664^7.36^, in the three different structures, assumes different conformations where it is flipped toward the intracellular side in the AZD5462-bound active state and flipped toward the extracellular side in the relaxin-2-bound active state (**Figure 6E**). F668^7.40^ exhibited a slight conformational shift in the AZD5462-bound state which perfectly aligned its aromatic ring parallel to the phenoxy ring of AZD5462 to form a π-π interaction (**Figure 6F**). We then tested the effect of these residues on AZD5462 binding using radioligand binding assay and found that the binding of AZD5462 to RXFP1 was completely abolished in W664A and F668A mutants (**Figure S6A**), confirming that these residues are the key determinants of AZD5462 binding to RXFP1. On the other hand, alanine mutation of Q408^1.35^, previously identified as an important residue and interacting partner of W664^7.36^ in the relaxin-2-mediated activation of RXFP1, did not affect binding affinity of AZD5462 (**Figure S6A**). Consistent with this result, no significant effect of this mutant was observed in the cAMP and β-arrestin recruitment assays in the AZD5462-mediated activation of RXFP1 (**Figures 6C and 6D**). Our functional data also indicated that point mutations of several residues surrounding the binding pocket that are involved in hydrophobic interactions with AZD5462, particularly S405A, V415A and I672A, resulted in decreased signaling of RXFP1, highlighting their importance in receptor activation (**Figure 6B; Table S7**).

Despite the different binding sites of the two agonists, it is clear that relaxin-2 and AZD5462 activation of RXFP1 involves shared residues (D637^6.44^, W664^>7.36^ and F668^7.40^) but distinct ligand-specific conformational changes. In the relaxin-2-bound active state, we observed an inward movement and rotation of TM7 that triggers activating signal to D637^6.44^ and W641^6.48^ of the transmission switch to induce receptor activation.^57^ However, in the AZD5462-bound state, this inward movement of the extracellular end of TM7 is absent, rather, the bulky leg of AZD5462 pushes the mid region of TM7 to cause an inward movement that clashes with D637^6.44^, thereby breaking the hydrogen bond interaction between W641^6.48^ and D637^6.44^, and consequently inducing the outward movement of TM6 to activate RXFP1 (**Figures 6H-6J**). Altogether, our data identifies a unique ligand-binding pocket of RXFP1, supports the binding mode of AZD5462 observed in our structure and identifies critical residues important for AZD5462 binding and activation of RXFP1.

## Discussion

RXFP1 is an emerging drug target, with at least three clinical efforts using both protein and small-molecule ligands in Phase 2 trials, but to date the molecular details of their actions remain unknown. In this study, we report the cryo-EM structures of RXFP1 inactive, relaxin-2-bound inactive and AZD5462-bound active states. Together with HDX, extensive mutagenesis and signaling assays, we define the mechanistic basis of relaxin-2 and AZD5462 recognition and activation of RXFP1. Relaxin-2 signaling requires an elaborate series of conformational relays to convert a ligand binding event into a conformational change. The flexible linker in the ECD adopts a defined secondary structure upon relaxin-2 binding. This conformational change consequently influences the physical location and interaction of the LDLa module to stabilize the active conformation of the receptor. This result explains prior data showing the essentiality of the linker and LDLa module in RXFP1 activation.^31–33^ Relaxin-2 binding in the distal ECD triggers conformational changes in the TMD to drive receptor activation. The hinge signaling motif of the ECD-TMD interface plays an essential role in transducing this signal from the ECD to the TMD. This finding is consistent with the regulatory role of the hinge in GPHRs, highlighting a universal signal transduction role of the hinge in the LGR subfamily.

In contrast, AZD5462 activates the receptor far more directly, binding to an unusual site in the TMD, in a manner more akin to a lipid than a conventional drug-like ligand. In doing so, it bypasses most of the structural mechanisms that relaxin-2 uses to activate signaling, including the requirement of the linker and LDLa module. Despite the different activation pathways, relaxin-2 and AZD5462 mechanisms engage a shared network of residues in TM7 to converge on activation of the transmission switch (CWxP motif) on TM6,^57^ directly resulting in the outward movement of TM6 for receptor activation. These findings illustrate a distinct activation mechanism from ligand binding to the transmission switch but a conserved pathway to G protein activation (**Figure 7**). However, AZD5462 also shows a unique signaling profile (β-arrestin recruitment), suggesting that it imperfectly recapitulates the features of relaxin-2. These results indicate that targeting other LGRs through corresponding sites may be possible, but attention to signaling details will be important. Soon, we will see how AZD5462 and protein drugs perform in the clinic, and it will be exciting to see if there are differential impacts from activating the receptor with a small molecule as compared to a protein agonist. Trials with relaxin-2 analogs such as TX45^26^ and AZD3427^25^ are currently ongoing, and the findings of these trails will shed more light on the clinical potential of long-acting human relaxin-2 in heart failure therapy.

**Figure 7.**
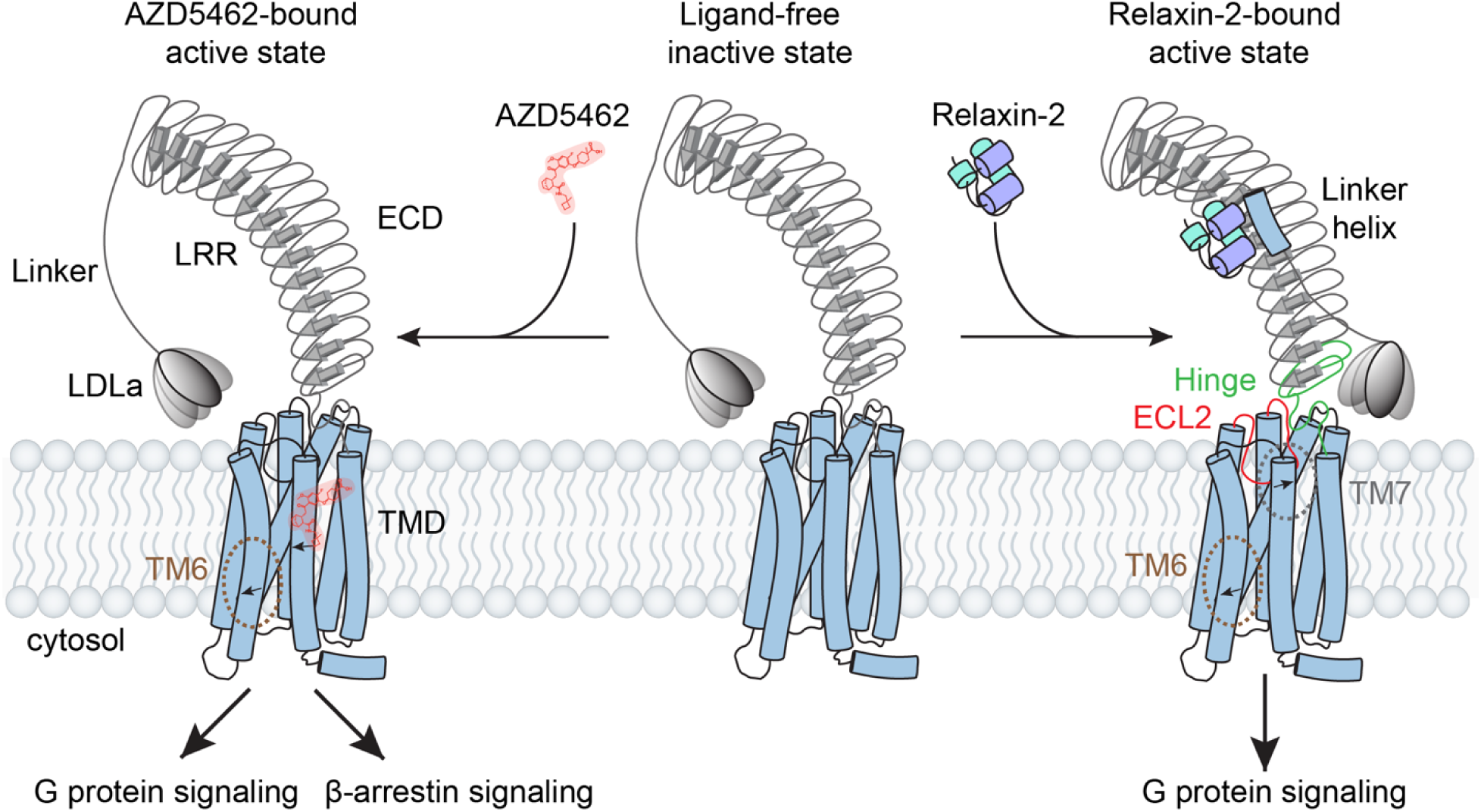
Schematic representation of model describing RXFP1 activation by Relaxin-2 and AZD5462. In the ligand-free (inactive) state, the linker and LDLa module are flexible. However, when relaxin-2 binds to the ECD, the flexible linker adopts a defined secondary structure. This conformational change consequently influences the orientation of the LDLa module to stabilize the active conformation of the receptor. Signal transduction from the ECD via the hinge region causes conformational changes in the TMD, particularly an inward movement of the upper region of TM7, increased hydrophobic contacts around the ECL2 and consequently an outward movement of TM6 to drive receptor activation to G protein signaling. Conversely, AZD5462 binds to an unusual site on the TMD, far from the relaxin-2 binding site and on the lipid-facing side of TM7. The bulky leg of AZD5462 pushes the mid region of TM7 to cause an inward movement that consequently induces the outward movement of TM6 to activate RXFP1 through the G protein and β-arrestin pathways.

Because AZD5462 and ML290 bind to the same ligand-binding pocket on the TMD of RXFP1, it would be intuitive to assume that AZD5462 and ML290 activate RXFP1 using the same underlying mechanism. However, our preliminary evidence together with previous data show that ML290 might activate RXFP1 using a different mechanism by engaging additional residues that do not significantly affect AZD5462-mediated activation of RXFP1. This is evident in the fact that AZD5462 can activate the F564A RXFP1 mutant whereas ML290 does not.^33^ Additionally, even though both AZD5462 and ML290-mediated signaling were abrogated in W664A and F668A mutants, only ML290-mediated activity was abolished in V415A whereas AZD5462 can still elicit receptor activation in this mutant (**Figure S6B**). Binding of ML290 to V415A was also impaired with a Ki of ∼143 nM, compared to ∼29 nM in wild type (WT) RXFP1 (**Figure S6C**). These data point to differences in the mechanisms of activation and might explain previously reported observations of differences in the signaling profile of ML290 and AZD5462, specifically that ML290 does not signal RXFP1 through the cGMP, ERK phosphorylation and β-arrestin recruitment pathways whereas AZD5462 signals through these pathways (**Figure S6D**).^29^ More work will be required to fully understand the details of ML290 activation of RXFP1.

Finally, despite the beneficial physiological properties of activating RXFP1 signaling, aberrant signaling leads to cancers of reproductive tissues, including endometrial, prostate and ovarian cancer.^11,12^ The availability of our RXFP1 structures therefore provides opportunities to rationally design and develop not only RXFP1 agonists but also antagonists for the treatment of RXFP1-associated cancers.

## Acknowledgements

Cryo-EM data was collected at the Harvard Center for Cryo-Electron Microscopy at Harvard Medical School, and we thank them for their support and advice during data collection. Structural biology applications used in this project were compiled and configured by SBGrid. We thank Marie Bao for critical reading of the manuscript. This work was funded by HHMI Hanna Gray fellows program awarded to J.O.-O., Blavatnik Biomedical Accelerator grant from Harvard Medical School to A.C.K., NIH R01 AR079489 to A.C.K. and M.W.G. and NIH Arthritis and Musculoskeletal and Skin Diseases K08AR084617 to J.S.S. The HDX-MS platforms used at the Montpellier Proteomics Platform (PPM, BioCampus) was co-financed by the European Regional Development Fund (ERDF) and the Occitanie region. PPM is a member of the national Proteomics French Infrastructure (ProFI UAR 2048) supported by the French National Research Agency (ANR-24-INBS-0015, Investments for the future F2030).

## Data availability

The cryo-EM maps and models were deposited under the following accession numbers: RXFP1 inactive: PDB-10RH, EMD-75403, relaxin-2-RXFP1 complex: PDB-10UZ, EMD-75473, AZD5462-RXFP1-miniGs complex: PDB-10SP, EMD-75439. Source data are provided with this paper.

## Author contributions

The molecular cloning, protein expression and purification, and cryo-EM grid preparation were performed by J.O.-O., X.W. and S.C.E. with supervision from A.C.K. Flow cytometry and cell signaling assays were performed by J.O.-O. and J.S.S with supervision by A.C.K. The cryo-EM data processing, model building, refinement and analyses were done by J.O.-O. with supervision from A.C.K and X.C. The HDX experiments and analysis were performed by T.G. with supervision from C.B. AZD5462 chemical synthesis was done by P.S. with supervision from D.E.K. The native relaxin-2 was provided by A.K.W. and M.W.G. The manuscript was written by J.O-O. and A.C.K. with input from all authors.

## Materials and Methods

### Cell lines

HEK293T cells were obtained from ATCC and were used for the GloSensor assay, BRET internalization assay, Fc-tagged relaxin-2 (SE301)^26^ binding experiments and determining surface expression levels of receptors. The cells were routinely maintained in Dulbecco’s modified Eagle’s medium (DMEM) (Corning) supplemented with 10% (v/v) fetal bovine serum (FBS) (Sigma-Aldrich) at 37°C in 5% CO_2_ incubator. HTLA cells were used for the PRESTO-TANGO assays and were maintained in DMEM supplemented with 10% FBS. Expi293F-TetR cells were used for receptor expression and were maintained in Expi293 media. Cell lines were regularly examined under the microscope for proper morphology and were also routinely tested for mycoplasma contamination.

### Cloning of RXFP1 constructs

The coding sequence of human RXFP1 from residues 23-757 (Uniprot ID Q9HBX9) with an N-terminal influenza hemagglutinin signal sequence and a FLAG tag (DYKDDDD) were cloned into a pcDNA-Zeo-tetO vector containing a tetracycline inducible cassette for all signaling assays and receptor expression determinations as previously described.^33^ All mutagenesis were introduced into plasmids using the QuikChange II XL site-directed mutagenesis kit (Agilent Technologies). For receptor expression and purification, we fused our construct with miniGs-414 with truncations of 15, 25 or 35 residues from the receptor’s C-terminus. To obtain inactive-state structure, we inserted a 3C proteolysis site between the receptor and the miniGs-414 using QuikChange II XL site-directed mutagenesis kit. This introduction, unexpectedly decreased receptor expression and hence to regain the expression level, we truncated RXFP1’s flexible C-terminus to identify the optimal expression construct (**Figures S1A and S1B**). This modified construct, having a C-terminal truncation of 30 amino acid residues, did not affect receptor signaling function. This was confirmed by cAMP accumulation assay showing a signaling response similar to that of the full receptor wild type when treated with relaxin-2 (**Figure S1E**). These modifications were made using PCR amplifications and NEBuilder HiFi DNA Assembly (New England Biolabs).

For β-arrestin recruitment, we cloned human RXFP1 from residues 23-757 with an N-terminal hemagglutinin signal sequence followed by a FLAG tag and a C-terminal TEV protease site followed by the tTA transcriptional activator into pcDNA-Zeo-tetO vector. The RXFP1 construct used in this assay was purposely engineered to exclude the C-terminus of the V2 vasopressin receptor (V2 tail) that promotes arrestin recruitment. This was to ensure that any signal of β-arrestin recruitment recorded could be attributed to RXFP1 only and not the V2 tail.

### Cell surface expression tests and flow cytometry binding assay

Cell surface expression levels were determined for RXFP1 signaling assay constructs as previously described.^33^ Briefly, HEK293T cells were plated at 100,000 cells/well into 12-well plates (Thermo Fisher Scientific). The cells were then transfected with RXFP1 WT or mutant signaling constructs using FuGENE HD (Promega), according to the manufacturer’s protocol (Promega). For each well, 200ng of construct was transfected unless stated otherwise and empty vector was transfected as negative control. To ensure that surface expression of WT was matched to mutants, serial dilutions of the WT construct were transfected. Twenty-four hours post transfection, plates were taken out of the incubator, the media was aspirated, and cells were washed with phosphate-buffered saline (PBS) with 1% (v/v) FBS and 2 mM calcium chloride (Buffer A). The cells were then detached by pipetting and distributed at 100,000 cells/well in 200 μL Buffer A into a V-bottom 96-well plate (Corning). The cells were blocked at 4°C for 20 min with shaking. This was followed by M1 anti-FLAG antibody (in house) labeled with Alexa Fluor 647 (Thermo Fisher Scientific) incubation for 30 min at 4°C with shaking. Cells were then washed once in Buffer A and resuspended in 100 μL Buffer A for flow cytometry analysis. Mean fluorescence intensity was quantified using a CytoFLEX flow cytometer (Beckman Coulter) and approximately 2,000 events per sample were collected. Data was normalized using the wild type and empty vector mean fluorescence intensities as 100% and 0%, respectively, and plotted using GraphPad Prism.

For flow cytometry binding assay, we used an Fc-tagged relaxin-2 (SE301) to measure binding. After blocking, cells were incubated with 500 nM SE301 for 1 h at 4°C with shaking. Cells were then washed twice and incubated with M1 anti-FLAG antibody (in house) labeled with Alexa Fluor 488 (in house) and Alexa Fluor-647 anti-human IgG Fc (BioLegend) for 30 min at 4°C with shaking. This was followed by washing and mean fluorescence intensity quantification using CytoFLEX flow cytometer.

### cAMP signaling assay

Relaxin receptor Gs activation and cAMP production were determined using the GloSensor cAMP assay (Promega) as previously described.^33^ In detail, white, clear-bottom 96-well plates (Thermo Fisher Scientific) were coated with 30 μL of 10 μg/mL poly-D-lysine (Sigma-Aldrich) for 30 min at room temperature. The plates were then washed twice with 100 μL PBS. HEK293T cells were seeded into the poly-D-lysine coated plates at 2×10^4^ cells/well. The cells were then co-transfected with 20 ng of GloSensor reporter plasmid and 20 ng (unless otherwise stated) of RXFP1, mutant or empty vector DNA per well using FuGENE HD (Promega), according to the manufacturer’s instructions. Twenty-four hours post-transfection, the media was aspirated and replaced with 40 μL of CO_2_-independent media (Thermo Fisher Scientific) with 10% (v/v) FBS and 2 mg/mL D-luciferin (Goldbio). The cells were then incubated for 2 h in the dark at room temperature. Luminescence was measured before and 30 min after ligand addition using a SpectraMax M5e microplate reader (Molecular Devices). Incubation was done in the dark at room temperature. Single concentration agonist-induced signaling were carried out using Emax concentrations, recombinant human relaxin-2 (R&D Systems) at 10 nM and AZD5462 (in house) at 1.25 µM. Data presented in Figures are normalized to % maximum response of wild type for each experiment and plotted using GraphPad Prism.

### β-arrestin-2 recruitment

The TANGO assay was used to determine the recruitment of β-arrestin-2 to RXFP1.^59^ HTLA cells stably expressing the tTA-dependent luciferase reporter and the β-arrestin-2-TEV protease fusion protein were seeded into 6-well plates. RXFP1 WT and mutant plasmids for β-arrestin-2 recruitment studies were transfected into the cells, 500 ng/well (unless otherwise stated), using FuGENE HD (Promega) according to the manufacturer’s protocol. Twenty-four hours post-transfection, cells were trypsinized and re-plated into poly-D-lysine coated 96-well clear bottom plates in serum- and phenol red-free DMEM. After 6 h, cells were stimulated with varying concentrations of AZD5462 or relaxin-2 or ML290 in a dose-dependent manner or at single point concentration for 14-16 h. An equal volume of Bright Glo buffer from the Bright-Glo Luciferase Assay System (Promega) was added to the cells for 10 min and luciferase expression was measured using a SpectraMax M5e plate reader (Molecular Devices). Data presented in Figures are normalized to % maximum response of wild type for each experiment and plotted using GraphPad Prism.

### Radioligand binding assay

Cell membranes for radioligand binding experiments were prepared from Expi293F TetR cells transiently transfected with either WT FLAG-RXFP1-3C-miniGs or mutant RXFP1 constructs. Transfection was done using FectoPro (400 μL/500 mL of culture) according to the manufacturer’s instructions. Cell cultures were enhanced with 3 mM valproic acid and 0.4% glucose 18 h after transfection. Receptor expression was then induced with 0.4 μg/mL doxycycline 24 h later. Cells were harvested 30 h after induction by centrifugation at 4,000 xg for 30 min at 4°C. The cell pellets were washed with cold HBS (20 mM HEPES pH 7.4, 150 mM NaCl) and resuspended in 2.5 ml of 20 mM Tris pH 7.4 per gram of cell pellet with protease inhibitor. The cells were then lysed by Dounce homogenization (approximately ×100). The cell membranes were pelleted by centrifugation at 50,000g for 20 min and resuspended in 2.5 ml of 50 mM Tris pH 7.4, 12.5 mM MgCl_2_, 150 mM NaCl, 0.2% BSA, and a protease inhibitor tablet by Dounce homogenization and flash-frozen in liquid nitrogen for storage at −80 °C.

For saturation curves, cell membranes of WT were incubated with varying concentrations of [^3^H]-AZD5462 (ViTrax Radiochemicals) in 50 mM Tris pH 7.4, 12.5 mM MgCl_2_, 150 mM NaCl, 0.2% BSA in a 200-μl reaction volume for 120 min at room temperature with shaking. Nonspecific binding of [^3^H]-AZD5462 was measured in the presence of 10 μM cold AZD5462. Cell membranes of WT or mutant RXFP1 were incubated with 2 nM [^3^H]-AZD5462 in 50 mM Tris pH 7.4, 12.5 mM MgCl_2_, 150 mM NaCl, 0.2% BSA in a 200-μl reaction volume for 120 min at room temperature with shaking. For competition binding assay, the membranes were incubated with 1 nM [^3^H]-AZD5462 and varying concentrations of ML290 in 50 mM Tris pH 7.4, 12.5 mM MgCl_2_, 150 mM NaCl, 0.2% BSA in a 200-μL reaction volume for 120 min at room temperature with shaking. Reactions were harvested on a GF/B filter soaked in water on a 96-well Brandel harvester and washed three times with cold water. Data were fit using a one-site saturation binding model (Total and non-specific) in GraphPad Prism to determine the radioligand affinity (K_D_). Data were fit using a one-site Ki to determine the inhibitory constant (Ki) values from competition binding data.

### RXFP1 BRET internalization assay

BRET internalization assay was conducted as previously described.^60^ Briefly, 200 ng of RXFP1 with a C-terminus NanoLuc tag expression plasmid, 400 ng myrpalm-Venus (to localize the dipole mVenus to the plasma membrane), 200 ng β-arrestin-1, and 200 ng β-arrestin-2 expression plasmids were transiently expressed in HEK293T cells seeded at ∼70% confluency in a six-well plate using FuGENE HD (Promega) according to manufacturer’s specifications. 18-24 h post transfection cells were swapped into a starvation medium (clear minimal essential media (MEM) supplemented with 1% penicillin and streptomycin, 1% anti-mycotic anti-biotic, 2% FBS) and seeded into a white, clear bottom, 96-well plate at 25,000-50,000 cells/well. After overnight incubation, starvation medium was removed and replaced with 90 μL of assay buffer containing 20 mM HEPES and 3 μM of coelenterazine-h diluted into Hanks Balanced Salt Solution buffer without calcium or magnesium (HBSS). BRET reads were conducted on a Promega plate reader with a 450 nM bandpass and a 530 nM longpass filter at 37°C as previously described.^61^ A BRET pre-read was conducted, cells were subsequently treated with 10 μM of AZD5462 or 100 nM, relaxin-2, and the plate was read multiple times for 20 min (final volume 100 μL). Pre-read BRET ratios were subtracted from post-read BRET ratios, and these values were normalized to vehicle treated wells to obtain the NET BRET ratio, with a greater negative value indicating RXFP1 movement away from the plasma membrane.

### Expression and Purification of RXFP1-miniGs414 and RXFP1-3C-miniGs414

RXFP1-miniGs or RXFP1-3C-miniGs plasmids (750 μg/L of culture) for protein expression and purification were transiently expressed in inducible Expi293F tetR cells (Thermo Fischer Scientific) using FectoPro (800 μL/L of culture) according to the manufacturer’s instructions. After 18 h, the culture was supplemented with 3 mM valproic acid and 0.4% glucose. Receptor expression was induced with 0.4 μg/mL doxycycline 48 hours post-transfection. Cells were harvested 30 hours after induction by centrifugation at 4,000 xg for 30 min at 4°C. The cell pellet was then flash frozen in liquid nitrogen and stored at −80°C until purification.

For receptor purification, frozen cells were hypotonically lysed in cold Lysis buffer containing 20 mM HEPES pH 7.5, 2 mM magnesium chloride, benzonase (Sigma Aldrich), protease inhibitor tablet (Thermo Fisher Scientific). For the relaxin-2-bound and AZD5462-bound states, 100 nM of relaxin-2 or 1000 nM of AZD5462 were added to all buffers throughout the purification. Iodoacetamide was added at a final concentration of 2 mg/ml during the cell lysis. The membrane fraction was collected by centrifugation at 50,000 xg for 30 min at 4°C. The membranes were then homogenized with a glass dounce homogenizer (Thermo Fisher Scientific) in solubilization buffer containing 20 mM HEPES pH 7.5, 350 mM sodium chloride, 20% (v/v) glycerol, 1% (w/v) lauryl maltose neopentyl glycol (L-MNG; Anatrace) and 0.1% (w/v) cholesterol hemisuccinate (CHS; Anatrace), benzonase, and protease inhibitor, and the solution was stirred for 2 h at 4°C to extract the receptor or ligand-receptor complex from the membrane. The soluble fraction was separated from the insoluble fraction by centrifugation at 50,000 xg for 30 min at 4°C and filtered with a glass fiber prefilter (Millipore). 2 mM Calcium chloride was added to the filtered solubilized fraction and incubated in homemade M1–Flag antibody-conjugated Sepharose resin by gravity flow at 4°C. The M1–sepharose resin was then washed extensively with wash buffer 1 [0.1% (w/v) L-MNG, 0.01% (w/v) CHS, 350 mM sodium chloride, 20 mM HEPES pH 7.5, 2 mM calcium chloride], wash buffer 2 [0.1% (w/v) L-MNG, 0.01% (w/v) CHS, 350 mM sodium chloride, 20 mM HEPES pH 7.5, 2 mM calcium chloride, 2 mM magnesium chloride, 5 mM adenosine 5’-triphosphate magnesium salt], and wash buffer 3 [0.01% (w/v) L-MNG, 0.001% (w/v) CHS, 350 mM sodium chloride, 20 mM HEPES pH 7.5, 2 mM calcium chloride] prior to elution with 0.01% (w/v) L-MNG, 0.001% (w/v) CHS, 350 mM sodium chloride, 20 mM HEPES pH 7.5, 0.5 mg/mL FLAG peptide (GenScript). The eluted protein was concentrated in a 100 kDa MWCO Amicon spin concentrator and injected onto a Superdex S200 Increase 10/300 column (GE Healthcare) equilibrated in size exclusion chromatography buffer [0.005% (w/v) L-MNG, 0.0005% (w/v) CHS, 350 mM sodium chloride, 20 mM HEPES pH 7.5]. For the RXFP1 inactive-state structure determination, the concentrated elute was incubated with 3C protease overnight at 4°C, prior to running on the Superdex S200 Increase 10/300 column. This was done to separate RXFP1 from the miniGs. The peak fractions from size exclusion were concentrated with a 3 kDa molecular weight cutoff centrifugal concentrator for preparation of cryo-EM grids or flash frozen in liquid nitrogen and stored at - 80°C.

### Cryo-EM grid preparation

Cryo-EM grids were prepared using an UltrAuFoil 300 mesh grids, 1.2-µm diameter/1.3-µm spacing (TedPella). The grids were glow-discharged at negative 15 mA for 30 seconds with a PELCO easiGlow. RXFP1, relaxin-2-RXFP1 or AZD5462-RXFP1-miniGs complex was concentrated to 0.5 mg/mL and 3 μL was applied to each grid. The grids were blotted for 4 to 6 s with a blot force of 15 s and wait time of 10 s on a Vitrobot Mark IV (ThermoFisher) at 10 °C and 100% humidity and then plunged into liquid-nitrogen-cooled ethane.

### Cryo-EM data collection and processing

Cryo-EM data for the inactive state RXFP1 and AZD5462-bound RXFP1-miniGs complex were collected in counted mode on a 300 kV Titan Krios G3i microscope (ThermoFisher) equipped with a K3 detector (Gatan) and GIF quantum energy Filter (20 eV) (Gatan) at the Harvard Cryo-Electron Microscopy Center for Structural Biology. The relaxin-2-bound RXFP1 data was collected on a Talos Arctica operating at 200 kV (ThermoFisher) equipped with K3 direct electron detector (Gatan) using Serial EM at Harvard Cryo-Electron Microscopy Center for Structural Biology.

Movies from the data sets were motion-corrected and dose-fractionated, followed by patch-based contrast transfer function (CTF) estimation, blob picking and local motion correction using CryoSPARC v3.^62^ For the ligand-free state RXFP1 dataset, initial 2D classification was used to generate templates which were used for template-based picking. The template-picked particles were subjected to multiple rounds of 2D classifications, followed an *ab initio* reconstruction. The resulting map was subjected to multiple rounds of heterogenous refinement, and the final stack of 226,677 particles was subjected to non-uniform refinement^63^ to a global resolution of 3.5 Å. The final map was further refined using DeepEMhancer.^64^

For the relaxin-2-bound RXFP1 dataset, initial 2D classification was used to generate templates which were used for template-based picking. The template-picked particles were subjected to multiple rounds of 2D classifications, followed an *ab initio* reconstruction. The resulting map was subjected to multiple rounds of heterogenous refinement, and the final stack of 325,329 particles was subjected to non-uniform refinement to a global resolution of 3.5 Å. The final map was further refined using DeepEMhancer.

For the AZD5462-bound RXFP1 dataset, initial 2D classification was used to generate templates which were used for template-based picking. The template-picked particles were subjected to multiple rounds of 2D classifications, followed an *ab initio* reconstruction. The resulting map was subjected to multiple rounds of heterogenous refinement, and the final stack of 69,698 particles was subjected to non-uniform refinement to a global resolution of 3.96 Å. The final map was further refined using DeepEMhancer. Data collection parameters are listed in Table S1.

### Model building and refinement

AlphaFold2^65^ predicted structure of RXFP1 was used as the starting model for building the ligand-free RXFP1 inactive model. A combination of the predicted structure and existing RXFP1-Gs-TM (accession no: 7TMW)^33^ structures were used to build the model for the AZD5462-bound RXFP1 structure. The miniGs of the AZD5462-bound RXFP1-miniGs complex was predicted using AlphaFold and was docked into the map separately, without additional model building. AlphaFold3^66^ predicted structure of RXFP1 was used as the starting model for building the relaxin-2-bound RXFP1 model. Using UCSF ChimeraX, the predicted structures were aligned and fitted into the experimental cryo-EM maps. The models were repeatedly refined with real space refinement in Phenix. The coordinates were further built manually in Coot.^67^ The structural biology softwares used in this project were compiled and configured by SBGrid software environment.^68^

### HDX-MS experiments

HDX-MS experiments were performed using a Synapt G2-Si HDMS coupled to nanoAQUITY UPLC with HDX Automation technology (Waters Corporation). RXFP1 in LMNG detergent was concentrated up to 20-25 µM and optimization of the sequence coverage was performed on undeuterated controls. The best sequence coverage and redundancy were obtained with Nepenthesin-2 protease (Affipro) with the addition of 2 M Urea and 200 or 400 mM TCEP in the quench buffer. Mixtures of receptor: ligands were pre-incubated together for 15 min at RT prior to HDX-MS analysis. Analysis of ligand-free RXFP1, RXFP1: AZD5462 (1: 1.6 or 30 ratio) and RXFP1: Relaxin-2 (1: 1.2 or 2 ratio) mixtures were performed as follows: 3 µL of sample are diluted in 57 µL of undeuterated for the reference or deuterated equilibration buffer (see HDX Summary Tables for buffer details). The final percentage of deuterium in the deuterated buffer was 95%. Deuteration was performed at 20°C for 0.5, 5, 15, 30 and 120 min. Next, 50 µL of reaction sample were quenched in 50 µL of quench buffer (KH_2_PO_4_ 50 mM, K_2_HPO_4_ 50 mM, 2 M Urea, 200 mM or 400 mM TCEP pH 2.3) at 0°C. 80 µL of quenched sample were loaded onto a 50 µL loop and injected on a Nepenthesin-2 column (Affipro), with 0.2% formic acid at a flowrate of 100 µL/min. The peptides were then trapped at 0°C on a Vanguard column (ACQUITY UPLC BEH C18 VanGuard Pre-column, 130 Å, 1.7 µm, 2.1 mm X 5 mm, Waters) for 3 min, before being loaded at 40 µL/min onto an Acquity UPLC column (ACQUITY UPLC BEH C18 Column, 1.7 µm, 1 mm X 100 mm, Waters) kept at 0°C. Peptides were subsequently eluted with a linear gradient (0.2% formic acid in acetonitrile solvent at 5% up to 35% during the first 6 min, then up to 40% and 95% over 1 min each) and ionized directly by electrospray on a Synapt G2-Si mass spectrometer (Waters). HDMSE data were obtained by 20-30 V trap collision energy ramp. Lock mass accuracy correction was made using a mixture of leucine enkephalin and GFP. For every tested condition we analyzed two biological replicates, representing different protein productions in expi293 mammalian cells, and deuteration timepoints were performed in triplicates for each condition.

Peptide identification was performed from undeuterated data using ProteinLynx global Server (PLGS, version 3.0.3, Waters). Peptides were filtered by DynamX (version 3.0, Waters) using the following parameters: minimum intensity of 1000, minimum product per amino acid of 0.12, maximum error for threshold of 10 ppm, and presence of peptides in at least 3 out of 6 files. All peptides were manually checked, and data was curated using DynamX. Back exchange was not corrected since we are measuring differential HDX and not absolute one. Statistical analysis of all ΔHDX data was performed using Deuteros 2.0^69^ and only peptides with a 99% confidence interval were considered.

## References

1 Petrie, E. J., Lagaida, S., Sethi, A., Bathgate, R. A. & Gooley, P. R. In a Class of Their Own - RXFP1 and RXFP2 are Unique Members of the LGR Family. Front Endocrinol (Lausanne) 6, 137 (2015). 10.3389/fendo.2015.00137

2 Scott, D. J. et al. Characterization of novel splice variants of LGR7 and LGR8 reveals that receptor signaling is mediated by their unique low density lipoprotein class A modules. J Biol Chem 281, 34942–34954 (2006). 10.1074/jbc.M602728200

3 Kong, R. C. et al. The relaxin receptor (RXFP1) utilizes hydrophobic moieties on a signaling surface of its N-terminal low density lipoprotein class A module to mediate receptor activation. J Biol Chem 288, 28138–28151 (2013). 10.1074/jbc.M113.499640

4 Hsu, S. Y. et al. Activation of orphan receptors by the hormone relaxin. Science 295, 671–674 (2002). 10.1126/science.1065654

5 MacLennan, A. H., Nicolson, R. & Green, R. C. Serum relaxin in pregnancy. Lancet 2, 241–243 (1986). 10.1016/s0140-6736(86)92068-4

6 Shabanpoor, F., Separovic, F. & Wade, J. D. The human insulin superfamily of polypeptide hormones. Vitam Horm 80, 1–31 (2009). 10.1016/S0083-6729(08)00601-8

7 Devarakonda, T. & Salloum, F. N. Heart Disease and Relaxin: New Actions for an Old Hormone. Trends Endocrinol Metab 29, 338–348 (2018). 10.1016/j.tem.2018.02.008

8 Valle Raleigh, J., et al. Reperfusion therapy with recombinant human relaxin-2 (Serelaxin) attenuates myocardial infarct size and NLRP3 inflammasome following ischemia/reperfusion injury via eNOS-dependent mechanism. Cardiovasc Res 113, 609–619 (2017). 10.1093/cvr/cvw246

9 Bennett, R. G. Relaxin and its role in the development and treatment of fibrosis. Transl Res 154, 1–6 (2009). 10.1016/j.trsl.2009.03.007

10 Fue, M. et al. Relaxin 2/RXFP1 Signaling Induces Cell Invasion via the β-Catenin Pathway in Endometrial Cancer. Int J Mol Sci 19 (2018). 10.3390/ijms19082438

11 Thanasupawat, T. et al. Emerging roles for the relaxin/RXFP1 system in cancer therapy. Mol Cell Endocrinol 487, 85–93 (2019). 10.1016/j.mce.2019.02.001

12 Burston, H. E. et al. Inhibition of relaxin autocrine signaling confers therapeutic vulnerability in ovarian cancer. J Clin Invest 131 (2021). 10.1172/JCI142677

13 Kirsch, J. R. et al. Minimally invasive, sustained-release relaxin-2 microparticles reverse arthrofibrosis. Sci Transl Med 14, eabo3357 (2022). 10.1126/scitranslmed.abo3357

14 Samuel, C. S. et al. Anti-fibrotic actions of relaxin. Br J Pharmacol 174, 962–976 (2017). 10.1111/bph.13529

15 Samuel, C. S. et al. Relaxin remodels fibrotic healing following myocardial infarction. Lab Invest 91, 675–690 (2011). 10.1038/labinvest.2010.198

16 Sarwar, M., Du, X. J., Dschietzig, T. B. & Summers, R. J. The actions of relaxin on the human cardiovascular system. Br J Pharmacol 174, 933–949 (2017). 10.1111/bph.13523

17 Samuel, C. S. et al. Relaxin deficiency in mice is associated with an age-related progression of pulmonary fibrosis. FASEB J 17, 121–123 (2003). 10.1096/fj.02-0449fje

18 Samuel, C. S. et al. The relaxin gene-knockout mouse: a model of progressive fibrosis. Ann N Y Acad Sci 1041, 173–181 (2005). 10.1196/annals.1282.025

19 Blessing, W. A. et al. Intraarticular injection of relaxin-2 alleviates shoulder arthrofibrosis. Proc Natl Acad Sci U S A 116, 12183–12192 (2019). 10.1073/pnas.1900355116

20 Teerlink, J. R. et al. Serelaxin, recombinant human relaxin-2, for treatment of acute heart failure (RELAX-AHF): a randomised, placebo-controlled trial. Lancet 381, 29–39 (2013). 10.1016/S0140-6736(12)61855-8

21 Teerlink, J. R. et al. Relaxin for the treatment of patients with acute heart failure (Pre-RELAX-AHF): a multicentre, randomised, placebo-controlled, parallel-group, dose-finding phase IIb study. Lancet 373, 1429–1439 (2009). 10.1016/S0140-6736(09)60622-X

22 Teichman, S. L. et al. Relaxin, a pleiotropic vasodilator for the treatment of heart failure. Heart Fail Rev 14, 321–329 (2009). 10.1007/s10741-008-9129-3

23 Metra, M. et al. Effects of Serelaxin in Patients with Acute Heart Failure. N Engl J Med 381, 716–726 (2019). 10.1056/NEJMoa1801291

24 Teerlink, J. R. et al. Effects of serelaxin in patients admitted for acute heart failure: a meta-analysis. Eur J Heart Fail 22, 315–329 (2020). 10.1002/ejhf.1692

25 Papworth, M. et al. A novel long-acting relaxin-2 fusion, AZD3427, improves cardiac performance in non-human primates with cardiac dysfunction. Cardiovasc Res 121, 871–881 (2025). 10.1093/cvr/cvaf031

26 Erlandson, S. C. et al. Engineering and Characterization of a Long-Half-Life Relaxin Receptor RXFP1 Agonist. Mol Pharm 21, 4441–4449 (2024). 10.1021/acs.molpharmaceut.4c00368

27 Borlaug, B. A. et al. Effects of volenrelaxin in worsening heart failure with preserved ejection fraction: a phase 2 randomized trial. Nat Med (2025). 10.1038/s41591-025-03939-6

28 McBride, A. et al. In search of a small molecule agonist of the relaxin receptor RXFP1 for the treatment of liver fibrosis. Sci Rep 7, 10806 (2017). 10.1038/s41598-017-10521-9

29 Granberg, K. L. et al. Discovery of Clinical Candidate AZD5462, a Selective Oral Allosteric RXFP1 Agonist for Treatment of Heart Failure. J Med Chem 67, 4419–4441 (2024). 10.1021/acs.jmedchem.3c02184

30 Medicine, N. L. o. (ClinicalTrials.gov, ClinicalTrials.gov, 2025).

31 Hoare, B. L. et al. Multi-Component Mechanism of H2 Relaxin Binding to RXFP1 through NanoBRET Kinetic Analysis. iScience 11, 93–113 (2019). 10.1016/j.isci.2018.12.004

32 Sethi, A. et al. The complex binding mode of the peptide hormone H2 relaxin to its receptor RXFP1. Nat Commun 7, 11344 (2016). 10.1038/ncomms11344

33 Erlandson, S. C. et al. The relaxin receptor RXFP1 signals through a mechanism of autoinhibition. Nat Chem Biol 19, 1013–1021 (2023). 10.1038/s41589-023-01321-6

34 Persechino, M., Hedderich, J. B., Kolb, P. & Hilger, D. Allosteric modulation of GPCRs: From structural insights to in silico drug discovery. Pharmacol Ther 237, 108242 (2022). 10.1016/j.pharmthera.2022.108242

35 Nehmé, R. et al. Mini-G proteins: Novel tools for studying GPCRs in their active conformation. PLoS One 12, e0175642 (2017). 10.1371/journal.pone.0175642

36 Carpenter, B. & Tate, C. G. Engineering a minimal G protein to facilitate crystallisation of G protein-coupled receptors in their active conformation. Protein Eng Des Sel 29, 583–594 (2016). 10.1093/protein/gzw049

37 Zhang, M. et al. G protein-coupled receptors (GPCRs): advances in structures, mechanisms, and drug discovery. Signal Transduct Target Ther 9, 88 (2024). 10.1038/s41392-024-01803-6

38 Duan, J. et al. Structures of full-length glycoprotein hormone receptor signalling complexes. Nature 598, 688–692 (2021). 10.1038/s41586-021-03924-2

39 Duan, J. et al. Hormone- and antibody-mediated activation of the thyrotropin receptor. Nature 609, 854–859 (2022). 10.1038/s41586-022-05173-3

40 Duan, J. et al. Mechanism of hormone and allosteric agonist mediated activation of follicle stimulating hormone receptor. Nat Commun 14, 519 (2023). 10.1038/s41467-023-36170-3

41 Diepenhorst, N. A. et al. Investigation of interactions at the extracellular loops of the relaxin family peptide receptor 1 (RXFP1). J Biol Chem 289, 34938–34952 (2014). 10.1074/jbc.M114.600882

42 Büllesbach, E. E. & Schwabe, C. The trap-like relaxin-binding site of the leucine-rich G-protein-coupled receptor 7. J Biol Chem 280, 14051–14056 (2005). 10.1074/jbc.M500030200

43 Scott, D. J., Tregear, G. W. & Bathgate, R. A. Modeling the primary hormone-binding site of RXFP1 and RXFP2. Ann N Y Acad Sci 1160, 74–77 (2009). 10.1111/j.1749-6632.2009.03950.x

44 Büllesbach, E. E., Yang, S. & Schwabe, C. The receptor-binding site of human relaxin II. A dual prong-binding mechanism. J Biol Chem 267, 22957–22960 (1992).

45 Büllesbach, E. E. & Schwabe, C. The relaxin receptor-binding site geometry suggests a novel gripping mode of interaction. J Biol Chem 275, 35276–35280 (2000). 10.1074/jbc.M005728200

46 Hossain, M. A. et al. The chemically synthesized human relaxin-2 analog, B-R13/17K H2, is an RXFP1 antagonist. Amino Acids 39, 409–416 (2010). 10.1007/s00726-009-0454-1

47 Hossain, M. A., Wade, J. D. & Bathgate, R. A. Chimeric relaxin peptides highlight the role of the A-chain in the function of H2 relaxin. Peptides 35, 102–106 (2012). 10.1016/j.peptides.2012.02.021

48 Bruell, S. et al. Chimeric RXFP1 and RXFP2 Receptors Highlight the Similar Mechanism of Activation Utilizing Their N-Terminal Low-Density Lipoprotein Class A Modules. Front Endocrinol (Lausanne) 4, 171 (2013). 10.3389/fendo.2013.00171

49 Chan, L. J. et al. Identification of key residues essential for the structural fold and receptor selectivity within the A-chain of human gene-2 (H2) relaxin. J Biol Chem 287, 41152–41164 (2012). 10.1074/jbc.M112.409284

50 Faust, B. et al. Autoantibody mimicry of hormone action at the thyrotropin receptor. Nature 609, 846–853 (2022). 10.1038/s41586-022-05159-1

51 Roos, P., Nyberg, L., Wide, L. & Gemzell, C. Human pituitary luteinizing hormone. Isolation and characterization of four glycoproteins with luteinizing activity. Biochim Biophys Acta 405, 363–379 (1975).

52 Carlsen, R. B., Bahl, O. P. & Swaminathan, N. Human chorionic gonadotropin. Linear amino acid sequence of the beta subunit. J Biol Chem 248, 6810–6827 (1973).

53 Gray, C. J. Molecular weight of human follicle stimulating hormone. Nature 216, 1112–1113 (1967). 10.1038/2161112a0

54 Wilson, K. J. et al. Optimization of the first small-molecule relaxin/insulin-like family peptide receptor (RXFP1) agonists: Activation results in an antifibrotic gene expression profile. Eur J Med Chem 156, 79–92 (2018). 10.1016/j.ejmech.2018.06.008

55 Xiao, J. et al. Identification and optimization of small-molecule agonists of the human relaxin hormone receptor RXFP1. Nat Commun 4, 1953 (2013). 10.1038/ncomms2953

56 Callander, G. E., Thomas, W. G. & Bathgate, R. A. Prolonged RXFP1 and RXFP2 signaling can be explained by poor internalization and a lack of beta-arrestin recruitment. Am J Physiol Cell Physiol 296, C1058–1066 (2009). 10.1152/ajpcell.00581.2008

57 Zhou, Q. et al. Common activation mechanism of class A GPCRs. Elife 8 (2019). 10.7554/eLife.50279

58 Laskowski, R. A. & Swindells, M. B. LigPlot+: multiple ligand-protein interaction diagrams for drug discovery. J Chem Inf Model 51, 2778–2786 (2011). 10.1021/ci200227u

59 Kroeze, W. K. et al. Author Correction: PRESTO-Tango as an open-source resource for interrogation of the druggable human GPCRome. Nat Struct Mol Biol 31, 578 (2024). 10.1038/s41594-023-01129-x

60 Smith, J. S. et al. C-X-C Motif Chemokine Receptor 3 Splice Variants Differentially Activate Beta-Arrestins to Regulate Downstream Signaling Pathways. Mol Pharmacol 92, 136–150 (2017). 10.1124/mol.117.108522

61 Smith, J. S. et al. The M3 Muscarinic Acetylcholine Receptor Can Signal through Multiple G Protein Families. Mol Pharmacol 105, 386–394 (2024). 10.1124/molpharm.123.000818

62 Punjani, A., Rubinstein, J. L., Fleet, D. J. & Brubaker, M. A. cryoSPARC: algorithms for rapid unsupervised cryo-EM structure determination. Nat Methods 14, 290–296 (2017). 10.1038/nmeth.4169

63 Punjani, A., Zhang, H. & Fleet, D. J. Non-uniform refinement: adaptive regularization improves single-particle cryo-EM reconstruction. Nat Methods 17, 1214–1221 (2020). 10.1038/s41592-020-00990-8

64 Sanchez-Garcia, R. et al. DeepEMhancer: a deep learning solution for cryo-EM volume post-processing. Commun Biol 4, 874 (2021). 10.1038/s42003-021-02399-1

65 Jumper, J. et al. Highly accurate protein structure prediction with AlphaFold. Nature 596, 583–589 (2021). 10.1038/s41586-021-03819-2

66 Abramson, J. et al. Accurate structure prediction of biomolecular interactions with AlphaFold 3. Nature 630, 493–500 (2024). 10.1038/s41586-024-07487-w

67 Emsley, P. & Cowtan, K. Coot: model-building tools for molecular graphics. Acta Crystallogr D Biol Crystallogr 60, 2126–2132 (2004). 10.1107/S0907444904019158

68 Herre, C. et al. Introduction of the Capsules environment to support further growth of the SBGrid structural biology software collection. Acta Crystallogr D Struct Biol 80, 439–450 (2024). 10.1107/S2059798324004881

69 Lau, A. M., Claesen, J., Hansen, K. & Politis, A. Deuteros 2.0: peptide-level significance testing of data from hydrogen deuterium exchange mass spectrometry. Bioinformatics 37, 270–272 (2021). 10.1093/bioinformatics/btaa677

