## Supplementary Information for "Divergent activation of the RXFP1 relaxin receptor by protein and small molecule agonists"

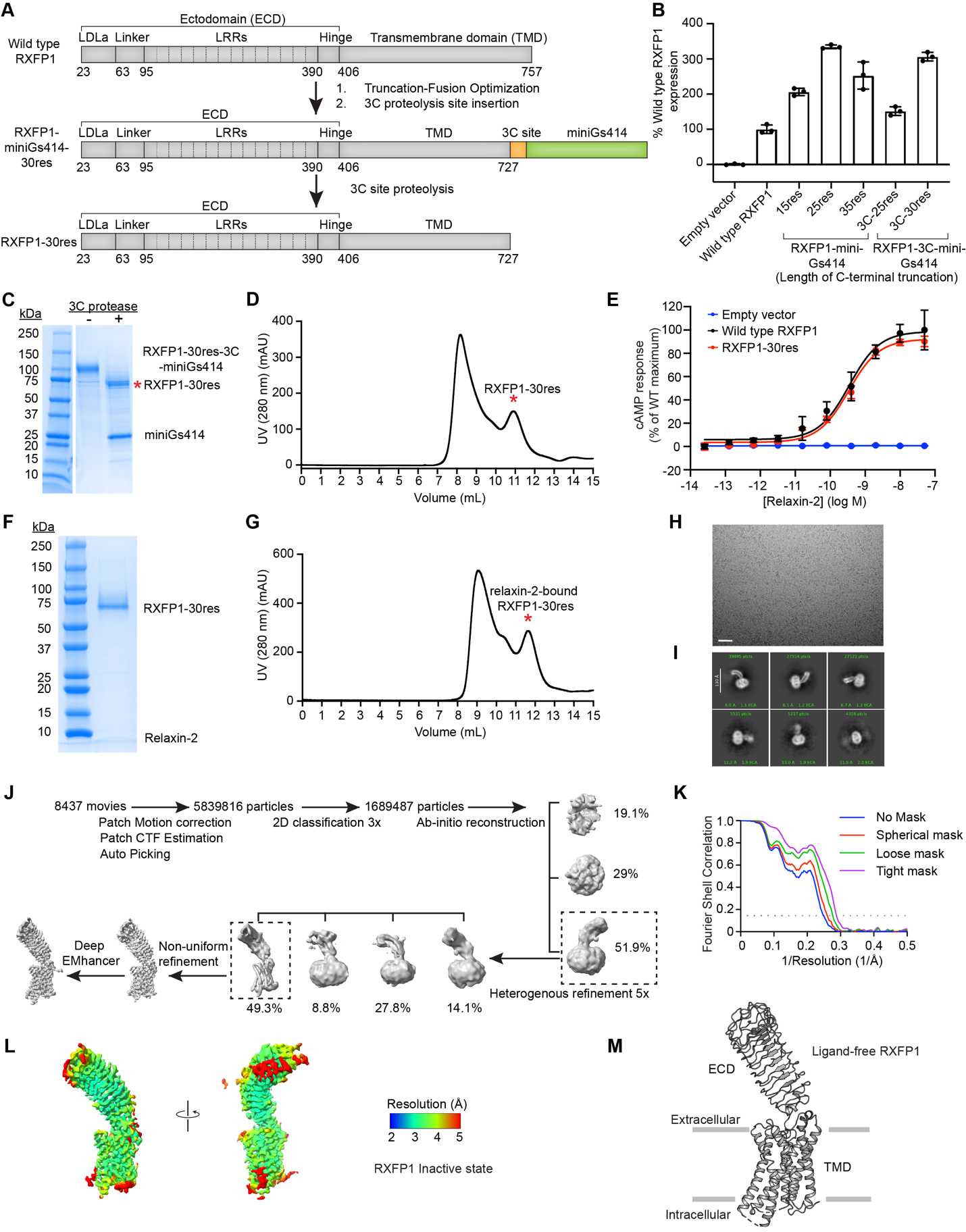

**Figure S1. Engineering, expression, purification, and structure determination of inactive-state RXFP1.** (**A**) Diagram of the primary structure of RXFP1 domains. Schematic illustrating the RXFP1 truncation, fusion, and 3C proteolysis site insertion. (**B**) Flow cytometry cell surface expression tests in Expi293F tetR cells for RXFP1-miniGs fusion constructs. Data is mean ± s.e.m., n = 3 biological replicates. (**C**) Representative image of a coomassie-stained SDS-PAGE gel for ligand-free RXFP1. (**D**) Size-exclusion chromatography profile for ligand-free RXFP1. Red asterisk indicates the peak fractions pooled for RXFP1 structure determination. (**E**) cAMP concentration-response curves for wild type RXFP1 versus C terminus truncated RXFP1 (RXFP1-30res) in response relaxin-2. Data is mean ± s.e.m., n = 3 biological replicates. (**F**) Representative image of a coomassie-stained SDS-PAGE gel for relaxin-2-bound RXFP1. (**G**) Size-exclusion chromatography profile for relaxin-2-bound RXFP1. Red asterisk indicates the peak fractions pooled for RXFP1 structure determination. (**H**) Representative micrograph from the ligand-free RXFP1 datasets obtained from Titan Krios (Scale bar = 50 nm). (**I**) Representative images of two-dimensional class averages of ligand-free RXFP1. (**J**) Cryo-EM data processing scheme for the ligand-free RXFP1. Shown are representative processing steps for the collected dataset. **k,** Fourier shell correlation (FSC) used to determine the overall map resolution. (**L**) cryoSPARC non-uniform refinement map colored by local resolution. (**M**) Cryo-EM model of ligand-free RXFP1 (PDB ID: 10RH).

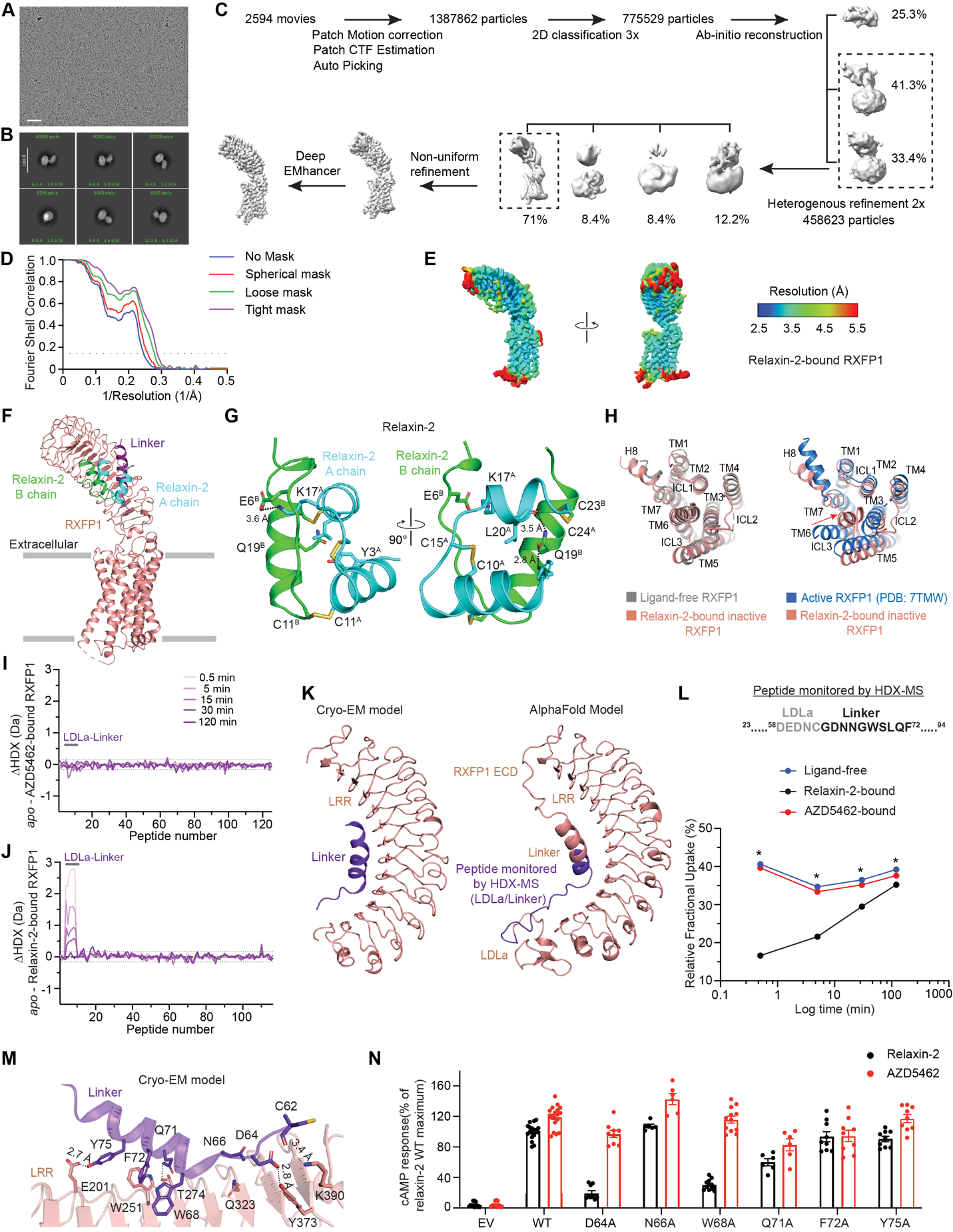

**Figure S2. Cryo-EM data processing for relaxin-2-bound RXFP1 and HDX-MS analysis.** (**A**) Representative micrograph from the relaxin-2-bound RXFP1 datasets obtained from a Talos Artica (Scale bar = 50 nm). (**B**) Representative images of two-dimensional class averages of relaxin-2-bound RXFP1. (**C**) Cryo-EM data processing scheme for the relaxin-2-bound RXFP1. Shown are representative processing steps for the collected dataset. (**D**) Fourier shell correlation (FSC) used to determine the overall map resolution. (**E**) cryoSPARC non-uniform refinement map colored by local resolution. (**F**) Cryo-EM model of relaxin-2-bound RXFP1 (PDB: 10UZ). (**G**) Cryo-EM model of relaxin-2, showing detailed interactions. (**H**) Conformational comparison of the intracellular view of (left) ligand-free inactive and relaxin-2-bound inactive RXFP1, (right) active and relaxin-2-bound inactive RXFP1. (**I** and **J**) Plots of the differences in the average deuterium uptake (ΔHDX) between different RXFP1 states for the identified peptides at five sampled time points. The individual peptides are arranged along the x-axis starting from the N- to the C-terminus. Positive and negative values indicate reduced and increased HDX, respectively. Values represent means of independent triplicate measurements. The dotted grey lines mark the threshold for a significant difference in HDX, which corresponds to the 99% confidence interval based on replicate measurements. ΔHDX of (I) RXFP1 in absence and presence of AZD5462 and (J) RXFP1 in absence and presence of Relaxin-2. (**K**) Structural representation of (left) RXFP1 LRR and linker in our cryo-EM model and (right) RXFP1 LRR, linker and LDLa module in AlphaFold model. Peptide monitored by HDX is illustrated in purple in the AlphaFold model. (**L**) Deuterium uptake plot showing the relative fractional uptake of the illustrated LDLa/Linker peptide in ligand-free (blue), relaxin-2-bound (black) and AZD5462-bound RXFP1 (red). The sequence of the corresponding residues is shown above. Uptake plots are the average and SD of 3 technical replicates from the same production of RXFP1. Statistically significant changes were determined using Deuteros 2.0 software (p ≤ 0.01). (**M**) Detailed interactions between the linker domain and the LRR. (**N**) Signaling in response to 10 nM relaxin-2 (black) or 1.25 μM AZD5462 (red) for RXFP1 linker mutants. Data are mean ± s.e.m. from three independent measurements.

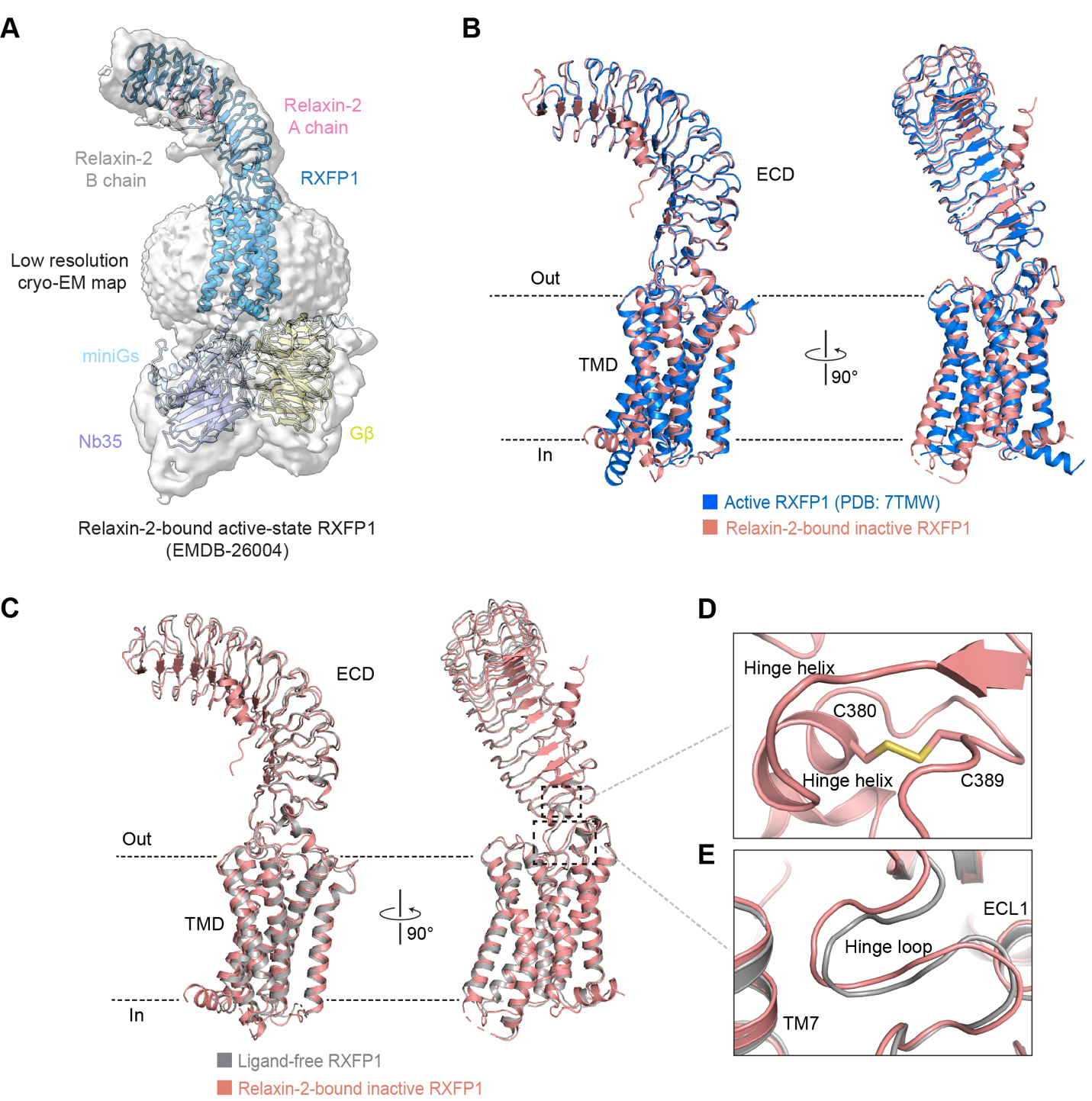

**Figure S3. Structural comparison of active and inactive RXFP1.** (**A**) RXFP1-miniGs complex model fit into low resolution cryo-EM map (EMDB-26004). Representative micrograph from relaxin-2-bound RXFP1 dataset. (**B**) Structural comparison of the active and relaxin-2-bound inactive RXFP1. (**C**) Structural comparison of the ligand-free inactive and relaxin-2-bound inactive RXFP1. (**D**) Detailed view highlighting the disulfide bond between C380 on the hinge helix and the C389 on the C terminal end of the LRR domain. (**E**) Conformational comparison of the hinge loop between ligand-free inactive and relaxin-2-bound inactive RXFP1.

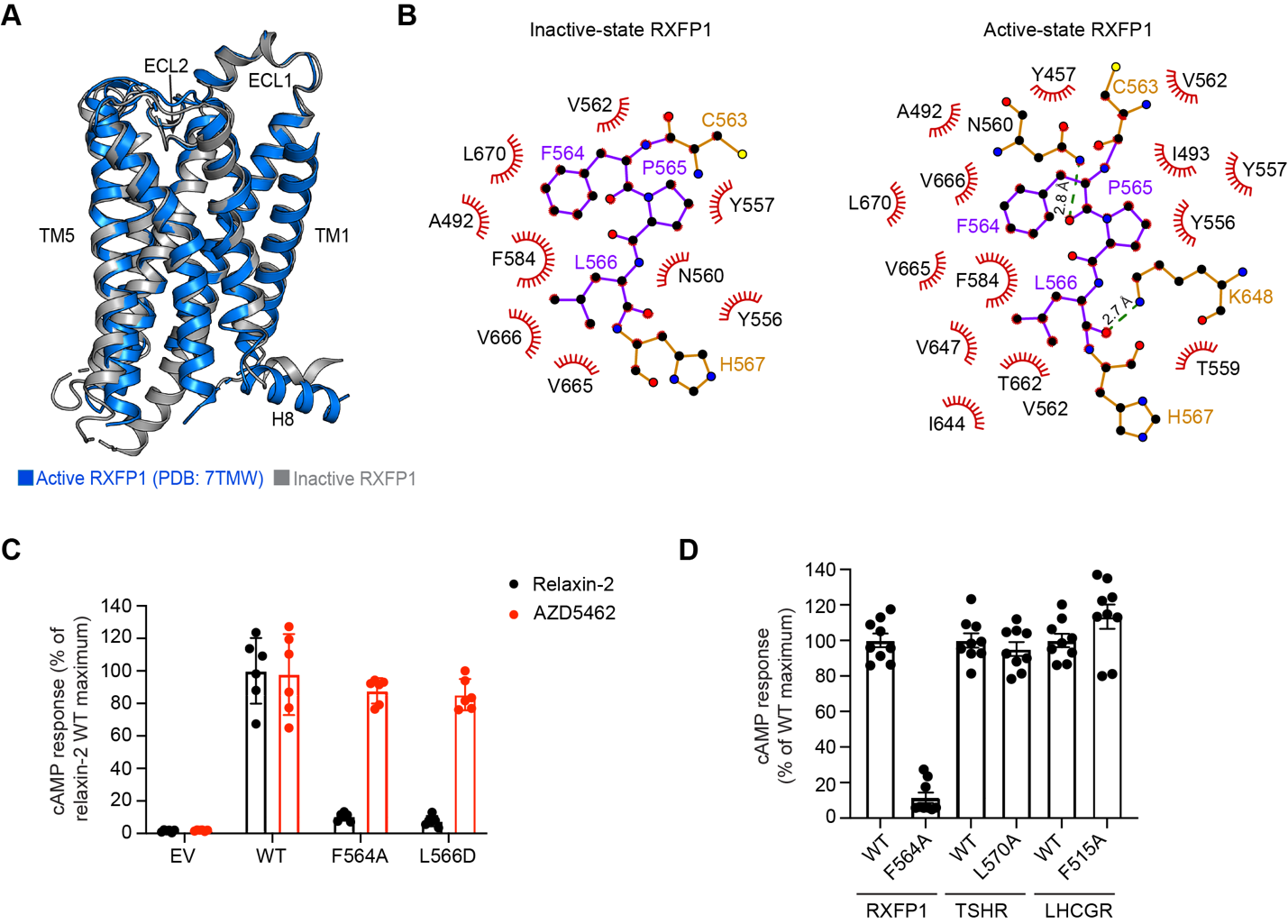

**Figure S4. Regulation of RXFP1 signaling by ECL2.** (**A**) Structural comparison of the active and inactive RXFP1. (**B**) Schematic representation of ECL2 interactions in the inactive versus active RXFP1. Generated using ligPlus program ECL2 residues are illustrated in purple. (**C**) Signaling in response to 10 nM relaxin-2 (black) or 1.25 μM AZD5462 (red) for RXFP1 ECL mutants. Data are mean ± s.e.m. from two independent experiments. (**D**) Signaling in response to relaxin-2 (10 nM), TSH (10 nM) and hCG (10 nM) for RXFP1, TSHR and LHCGR ECL2 mutants respectively. Data are mean ± s.e.m. from at least three independent experiments. Effect of relaxin-2 and AZD5462 on the role of ECL2 in RXFP1 activation.

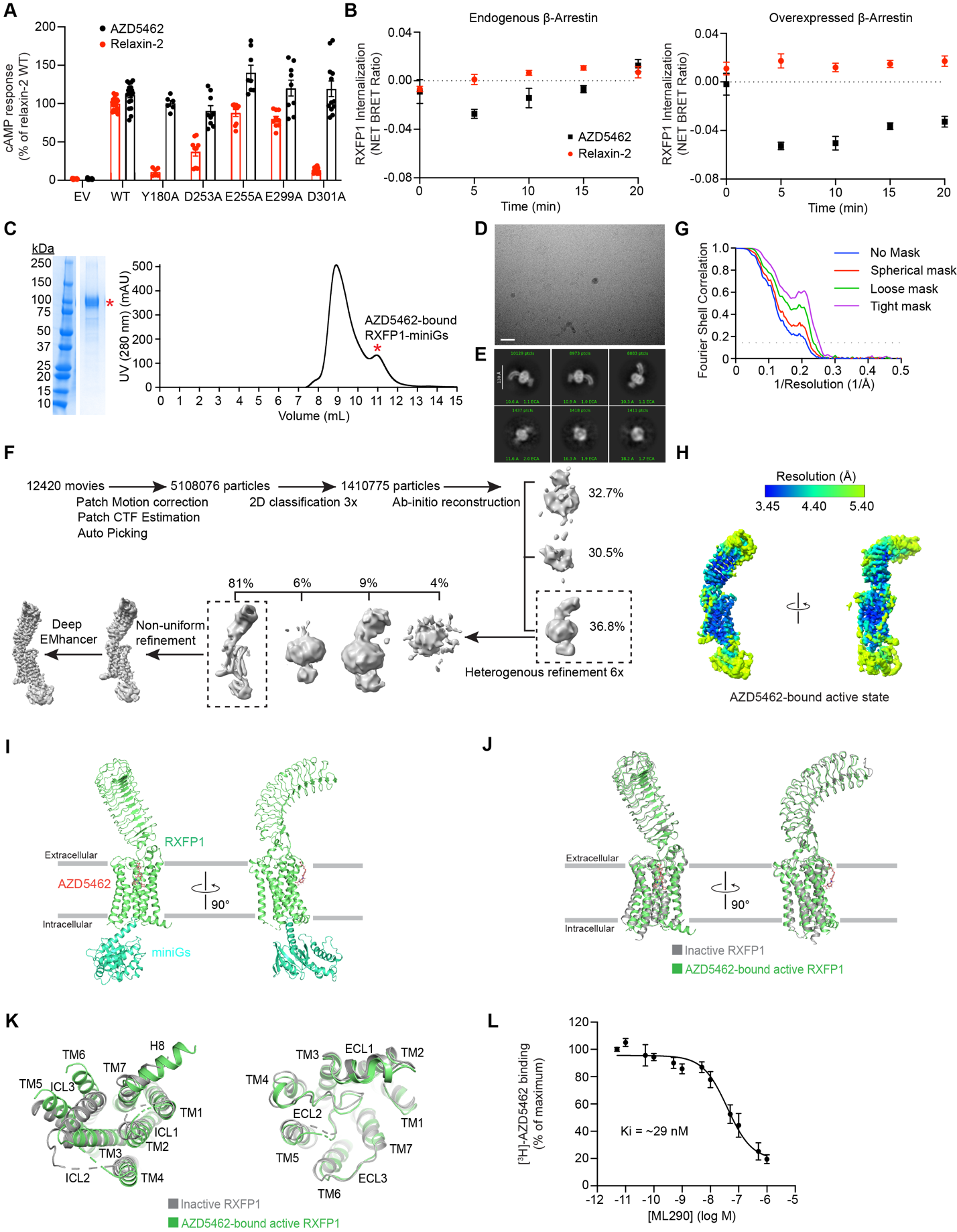

**Figure S5. Functional and structural characterization of AZD5462 activation of RXFP1.** (**A**) Signaling in response to 10 nM relaxin-2 (black) or 1.25 μM AZD5462 (red) for RXFP1 LRR mutants. Data are mean ± s.e.m. from at least three independent experiments. (**B**) AZD5462 (10 μM) or relaxin-2 (0.1 μM) -mediated RXFP1 internalization for 0, 5, 10, 15 or 20-min in endogenous and overexpressed β-arrestin systems. Pre-read BRET ratios were subtracted from post-read BRET ratios, and these values were normalized to vehicle treated wells to obtain the NET BRET ratio. Data are represented as the means ± s.e.m. of at least three independent experiments. (**C**) Representative image of a coomassie-stained SDS-PAGE gel for AZD5462-bound RXFP1-miniGs complex. Size-exclusion chromatography profile for AZD5462-bound RXFP1-miniGs complex. Red asterisk indicates the peak fractions pooled for RXFP1 structure determination. (**D**) Representative micrograph from the AZD5462-bound RXFP1-miniGs complex dataset obtained from a Titan Krios (Scale bar = 50 nm). (**E**) Representative images of two-dimensional class averages of AZD5462-bound RXFP1-miniGs complex. (**F**) Cryo-EM data processing scheme for the AZD5462-bound RXFP1-miniGs complex. Shown are representative processing steps for the collected dataset. (**G**) Fourier shell correlation (FSC) used to determine the overall map resolution. (**H**) cryoSPARC non-uniform refinement map colored by local resolution. (**I**) Cryo-EM model of AZD5462-bound RXFP1-miniGs complex (PDB ID: 10SP). (**J**) Structural comparison of the ligand-free inactive and AZD5462-bound active RXFP1. (**K**) Intracellular and extracellular view of the structural comparison of the ligand-free inactive and AZD5462-bound active RXFP1. (**L**) Radioligand competition binding experiment using AZD5462 (^3^H-AZD5462) and ML290. ML290 displaces ^3^H-AZD5462 from cell membranes containing wild type RXFP1. Data are presented as mean ± s.e.m. from two independent measurements.

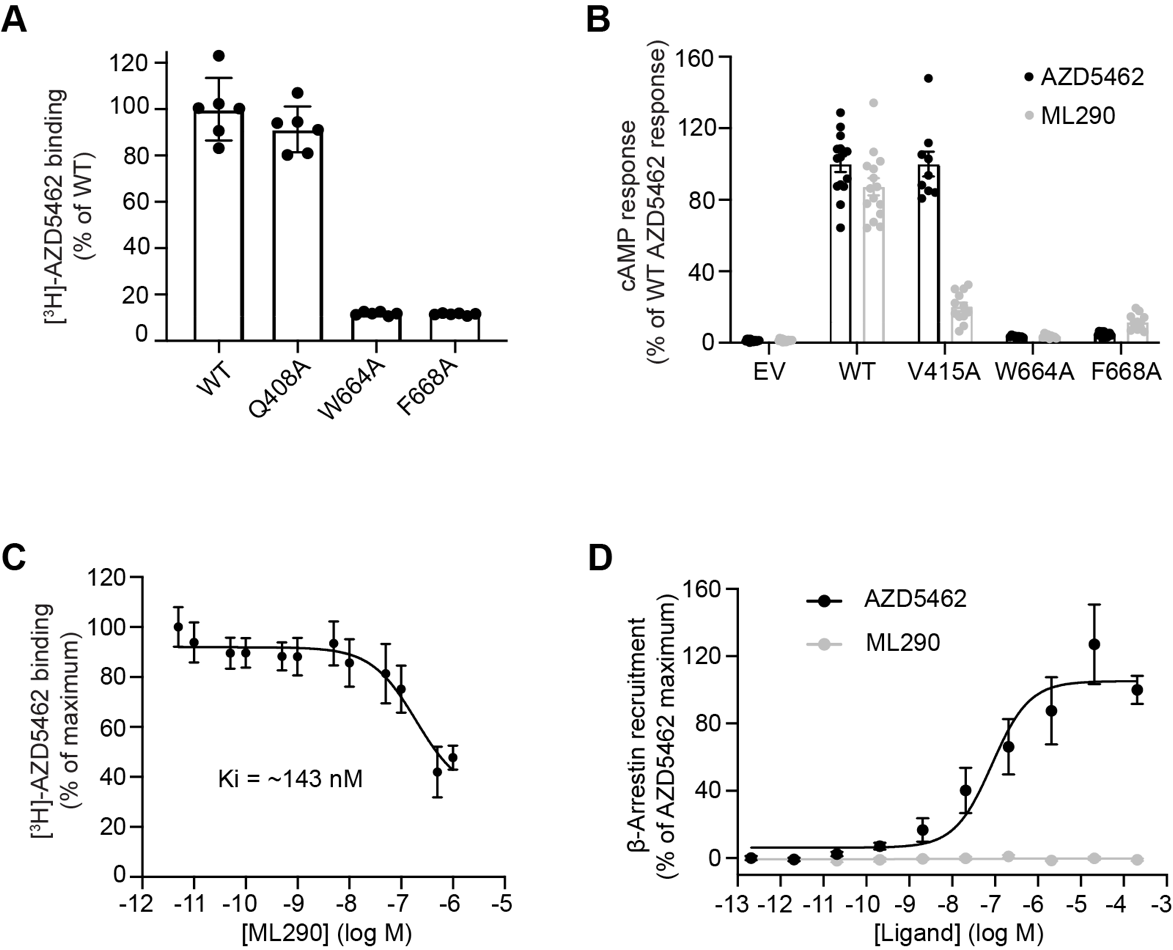

**Figure S6. Activation mechanism and signaling profile of RXFP1 by ML290.** (**A**) Total binding of radiolabeled AZD5462 (^3^H-AZD5462) to cell membranes containing wild type and mutant RXFP1. Data are mean ± s.e.m. from two independent experiments. (**B**) Signaling in response to 1.25 μM AZD5462 (black) or 60 μM ML290 (grey) for RXFP1 mutants. Data are mean ± s.e.m. from at least three independent experiments. (**C**) Radioligand competition binding experiment using AZD5462 (^3^H-AZD5462) and ML290. ML290 displaces ^3^H-AZD5462 from cell membranes containing V415A mutant. Data are presented as mean ± s.e.m. from two independent measurements. (**D**) β-Arrestin concentration-response curves for WT RXFP1 in response to AZD5462 (same as illustrated in Figure 5D) or ML290. Data are mean ± s.e.m. from at least three independent experiments.

**Table S1. Cryo-EM data collection, refinement, and validation statistics.**

|  | RXFP1 | Relaxin-2-RXFP1 | AZD5462-RXFP1-mGs |
| --- | --- | --- | --- |
|  | (PDB-10RH) (EMD-75403) | (PDB-10UZ) (EMD-75473) | (PDB-10SP) (EMD-75439) |
| Cryo-EM data collection and processing |  |  |  |
| Magnification | 105,000 | 36,000 | 105,000 |
| Voltage (kV) | 300 | 200 | 300 |
| Electron exposure (e-/ Å^2^) | ~63 | ~52 | 63 |
| Defocus range (μm) | -0.9 to -2.0 | -0.8 to -2.2 | -0.8 to -2.2 |
| Pixel size (Å) | 0.825 | 1.1 | 0.825 |
| Symmetry | C1 | C1 | C1 |
| Initial particle images (no.) | 5839816 | 1387862 | 5108076 |
| Final particle images (no.) | 226,677 | 325,329 | 69,698 |
| Map resolution (Å) | 3.5 | 3.5 | 3.96 |
| FSC threshold | (0.143) | (0.143) | 0.143 |
| Model refinement and validation |  |  |  |
| Initial model used (PDB) | 7TMW and AlphaFold2 | AlphaFold3 | 7TMW and AlphaFold2 |
| Map sharpening *B* factor | DeepEMhancer | DeepEMhancer | DeepEMhancer |
| Model composition |  |  |  |
| Non-hydrogen atoms | 4646 | 5382 | 5897 |
| Protein residues | 574 | 668 | 758 |
| Ligands | 0 | 1 | 1 |
| R.m.s. deviations |  |  |  |
| Bond lengths (Å) | 0.002 | 0.003 | 0.002 |
| Bond angles (Å) | 0.55 | 0.66 | 0.44 |
| Validation |  |  |  |
| MolProbity score | 1.97 | 2.1 | 3.12 |
| Clashscore | 10 | 8 | 51 |
| Poor rotamers (%) | 0.8 | 1.2 | 0.6 |
| Ramachandran plot |  |  |  |
| Favored (%) | 96 | 95 | 94 |
| Allowed (%) | 4 | 5 | 6 |
| Disallowed (%) | 0 | 0 | 0 |

**Table S2.** **Summary of Gs signaling, binding and expression data for RXFP1 constructs in Figure 2F.** Values are expressed as mean pEC_50_, E_max_, binding and cell surface expression ± s.e.m. from (n) biological replicates. ND = Not determined.

| Construct | pEC_50_ (Relaxin-2) | E_max_ (%) | Binding (%) | Cell surface expression (%) |
| --- | --- | --- | --- | --- |
| Wild type (WT) | 9.9 ± 0.1 (9) | 100 ± 4 (9) | 100 ± 3 (14) | 100 ± 1 (25) |
| Empty vector (EV) | ND | 0.3 ± 0.1 (9) | 0.6 ± 1 (14) | 0 ± 1.6 (21) |
| D253A | 8.2 ± 0.2 (9) | 67 ± 11 (9) | 13 ± 2 (12) | 75 ± 9 (13) |
| E255A | 9.2 ± 0.1 (9) | 105 ± 9 (9) | 50 ± 9 (8) | 113 ± 4 (16) |
| E299A | 8.5 ± 0.1 (9) | 81 ± 6 (9) | 51 ± 7 11) | 199 ± 11 (19) |
| D301A | Poor fit | 9 ± 1 (9) | 19 ± 4 (11) | 59 ± 4 (16) |

**Table S3.** **Summary of Gs signaling and expression data for RXFP1 constructs in Figures 2G and 4F.** Values are expressed as mean pEC_50_, E_max_ and cell surface expression ± s.e.m. from (n) biological replicates. ND = Not determined.

| Construct | pEC_50_ (Relaxin-2) | E_max_ (%) | Cell surface expression (%) |
| --- | --- | --- | --- |
| Wild type (WT) | 10 ± 0.1 (8) | 100 ± 5 (8) | 100 ± 3 (18) |
| Y180 | 7.5 ± 0.2 (8) | 23 ± 5 (8) | 21 ± 7 (3) |
| I204A | 9.5 ± 0.1 (8) | 92 ± 11 (8) | 41 ± 7 (9) |
| L226A | 9.3 ± 0.1 (8) | 123 ± 5 (8) | 70 ± 5 (9) |
| V228A | 9.8 ± 0.1 (8) | 107 ± 15 (8) | 95 ± 7 (15) |
| Wild type (WT) | 10 ± 0.1 (8) | 100 ± 4 (8) | 100 ± 5 (12) |
| D73A | 9.4 ± 0.1 (9) | 106 ± 9 (9) | ND |
| F76A | 9.8 ± 0.1 (6) | 89 ± 12 (6) | 420 ± 33 (3) |
| Y80A | 9.5 ± 0.1 (8) | 109 ± 6 (8) | 44 ± 6 (12) |
| Wild type (WT) | 10 ± 0.1 (6) | 100 ± 3 (6) | 100 ± 2 (21) |
| Empty vector (EV) | ND | 0.2 ± 0.2 (3) | 0 ± 0.7 (21) |
| Q408A | 9.0 ± 0.1 (6) | 67 ± 4 (6) | 99 ± 10 (12) |
| W664A | Poor fit | 14 ± 2 (6) | 85 ± 6 (15) |
| F668A | 9.0 ± 0.1 (6) | 52 ± 6 (6) | 67 ± 4 (15) |

**Table S4.** **Summary of Gs signaling and expression data for RXFP1 constructs in Figures S2N, S4C and 6G.** Values are expressed as mean E_max_ (relaxin-2 and AZD5462) and cell surface expression ± s.e.m. from (n) biological replicates.

| Construct | E_max_ (%) (Relaxin-2) | E_max_ (%) (AZD5462) | Cell surface expression (%) |
| --- | --- | --- | --- |
| Wild type (WT) | 100 ± 2 (12) | 119 ± 4 (12) | 100 ± 2 (9) |
| Empty vector (EV) | 4.5 ± 1 (9) | 4.7 ± 1 (9) | 0 ± 0.6 (3) |
| D64A | 19.7 ± 3 (9) | 97 ± 5 (9) | 72 ± 2 (3) |
| N66A | 107 ± 2 (6) | 143 ± 7 (6) | 93 ± 8 (3) |
| W68A | 30 ± 2 (12) | 116 ± 5 (12) | 84 ± 3 (3) |
| Q71A | 60 ± 5 (6) | 83 ± 8 (6) | 90 ± 7 (9) |
| F72A | 93 ± 7 (9) | 95 ± 8 (9) | 91 ± 8 (3) |
| Y75A | 90 ± 4 (9) | 117 ± 5 (9) | 113 ± 6 (9) |
| Wild type (WT) | 100 ± 8 (6) | 98 ± 10 (6) | 100 ± 3 (12) |
| Empty vector (EV) | 2 ± 0.3 (6) | 2 ± 0.2 (6) | 0 ± 1 (9) |
| F564A | 10 ± 1 (6) | 88 ± 3 (6) | 52 ± 2 (9) |
| L566D | 8 ± 1 (6) | 85 ± 4 (6) | 71 ± 10 (9) |
| Wild type (WT) | 100 ± 5 (9) | 122 ± 8 (9) | 100 ± 2 (21) |
| Empty vector (EV) | 1 ± 0.2 (9) | 1 ± 0.2 (9) | 0 ± 0.7 (21) |
| W664A | 4 ± 0.1 (3) | 5 ± 0.2 (3) | 85 ± 6 (15) |
| F668A | 64 ± 6 (9) | 7 ± 0.4 (9) | 67 ± 4 (15) |

**Table S5.** **Summary of Gs signaling for RXFP1 constructs in Figure 3C and 4G.** Values are expressed as mean E_max_ of basal and relaxin-2 signaling ± s.e.m. from (n) biological replicates.

| Construct | E_max_ (%) (Basal) | E_max_ (%) (Relaxin-2) |
| --- | --- | --- |
| Wild type (WT) | 11 ± 3 (6) | 100 ± 2 (6) |
| Empty vector (EV) | 1 ± 0.2 (3) | 1.4 ± 0.4 (3) |
| T393A | 78 ± 2 (3) | 108 ± 8 (3) |
| G395A | 63 ± 6 (3) | 70 ± 1 (3) |
| I396A | 45 ± 4 (3) | 66 ± 1 (3) |
| S397A | 70 ± 8 (3) | 95 ± 7 (3) |
| S398A | 54 ± 2 (3) | 77 ± 0.1 (3) |
| L399A | 106 ± 5 (3) | 98 ± 5 (3) |
| E400A | 1.5 ± 0.1 (3) | 90 ± 7 (3) |
| Wild type (WT) | 6 ± 1 (6) | 100 ± 2 (6) |
| Empty vector (EV) | 1 ± 0.1 (3) | 1.4 ± 0.4 (3) |
| D637A | 51 ± 6 (6) | 111 ± 3 (6) |

**Table S6.** **Summary of Gs signaling for RXFP1, TSHR and LHCGR constructs in Figure S4D.** Values are expressed as mean E_max_ and cell surface expression ± s.e.m. from (n) biological replicates.

| Construct | E_max_ (%) (Agonist) | Cell surface expression (%) |
| --- | --- | --- |
| WT RXFP1 | 100 ± 4 (9) | 100 ± 2 (6) |
| F564A | 11 ± 3 (9) | 77 ± 3 (6) |
| WT TSHR | 100 ± 4 (9) | 100 ± 13 (6) |
| L570A | 95 ± 4 (9) | 76 ± 11 (6) |
| WT LHCGR | 100 ± 4 (9) | 100 ± 4 (6) |
| F515A | 113 ± 7 (9) | 169 ± 25 (6) |

**Table S7.** **Summary of Gs signaling and expression data for RXFP1 constructs in Figure 6C.** Values are expressed as mean pEC_50_, E_max_ and cell surface expression ± s.e.m. from (n) biological replicates. ND = Not determined.

| Construct | pEC_50_ (AZD5462) | E_max_ (%) | Cell surface expression (%) |
| --- | --- | --- | --- |
| Wild type (WT) | 8.5 ± 0.1 (9) | 100 ± 2 (9) | 100 ± 2 (21) |
| Empty vector (EV) | ND | 0.3 ± 0.3 (9) | 0 ± 0.7 (21) |
| Q408A | 8.5 ± 0.1 (9) | 76 ± 7 (9) | 99 ± 10 (12) |
| W664A | 5.3 ± 0.3 (9) | 22 ± 6 (9) | 85 ± 6 (15) |
| F668A | 4.6 ± 0.9 (9) | 16 ± 3 (9) | 67 ± 4 (15) |
| P671A | 5.3 ± 0.4 (3) | 10 ± 3 (3) | 113 ± 11 (6) |
| Wild type (WT) | 8.1 ± 0.1 (12) | 100 ± 5 (12) | 100 ± 5 (12) |
| Empty vector (EV) | ND | 0.4 ± 0.2 (12) | ND |
| S405A | 6 ± 0.1 (9) | 96 ± 15 (12) | 72 ± 8 (6) |
| F411A | 8.4 ± 0.1 (8) | 90 ± 7 (12) | 81 ± 5 (3) |
| V415A | 7.5 ± 0.1 (12) | 126 ± 13 (12) | 85 ± 7 (9) |
| L639A | 8.6 ± 0.1 (12) | 117 ± 21 (12) | 74 ± 13 (6) |
| I672A | 6.6 ± 0.1 (12) | 87 ± 8 (12) | 71 ± 13 (6) |
